# Locally adaptive Bayesian birth-death model successfully detects slow and rapid rate shifts

**DOI:** 10.1101/853960

**Authors:** Andrew F. Magee, Sebastian Höhna, Tetyana I. Vasylyeva, Adam D. Leaché, Vladimir N. Minin

**Affiliations:** Department of Biology, University of Washington, Seattle, WA, 98195, USA; GeoBio-Center, Ludwig-Maximilians-Universität München, 80333 Munich, Germany; Department of Earth and Environmental Sciences, Paleontology & Geobiology, Ludwig-Maximilians-Universität München, 80333 Munich, Germany; Department of Zoology, University of Oxford, Oxford, United Kingdom; Department of Statistics, University of California, Irvine, CA, 92697, USA

## Abstract

Birth-death processes have given biologists a model-based framework to answer questions about changes in the birth and death rates of lineages in a phylogenetic tree. Therefore birth-death models are central to macroevolutionary as well as phylodynamic analyses. Early approaches to studying temporal variation in birth and death rates using birth-death models faced difficulties due to the restrictive choices of birth and death rate curves through time. Sufficiently flexible time-varying birth-death models are still lacking. We use a piecewise-constant birth-death model, combined with both Gaussian Markov random field (GMRF) and horseshoe Markov random field (HSMRF) prior distributions, to approximate arbitrary changes in birth rate through time. We implement these models in the widely used statistical phylogenetic software platform `RevBayes`, allowing us to jointly estimate birth-death process parameters, phylogeny, and nuisance parameters in a Bayesian framework. We test both GMRF-based and HSMRF-based models on a variety of simulated diversification scenarios, and then apply them to both a macroevolutionary and an epidemiological dataset. We find that both models are capable of inferring variable birth rates and correctly rejecting variable models in favor of effectively constant models. In general the HSMRF-based model has higher precision than its GMRF counterpart, with little to no loss of accuracy. Applied to a macroevolutionary dataset of the Australian gecko family Pygopodidae (where birth rates are interpretable as speciation rates), the GMRF-based model detects a slow decrease whereas the HSMRF-based model detects a rapid speciation-rate decrease in the last 12 million years. Applied to an infectious disease phylodynamic dataset of sequences from HIV subtype A in Russia and Ukraine (where birth rates are interpretable as the rate of accumulation of new infections), our models detect a strongly elevated rate of infection in the 1990s.

**Author summary:** Both the growth of groups of species and the spread of infectious diseases through populations can be modeled as birth-death processes. Birth events correspond either to speciation or infection, and death events to extinction or becoming noninfectious. The rates of birth and death may vary over time, and by examining this variation researchers can pinpoint important events in the history of life on Earth or in the course of an outbreak. Time-calibrated phylogenies track the relationships between a set of species (or infections) and the times of all speciation (or infection) events, and can thus be used to infer birth and death rates. We develop two phylogenetic birth-death models with the goal of discerning signal of rate variation from noise due to the stochastic nature of birth-death models. Using a variety of simulated datasets, we show that one of these models can accurately infer slow and rapid rate shifts without sacrificing precision. Using real data, we demonstrate that our new methodology can be used for simultaneous inference of phylogeny and rates through time.

## Introduction

Studying variation in the rates of speciation and extinction enables researchers to examine the patterns and processes that shape the diversity of life on earth. Birth-death processes have given biologists a model-based framework in which questions about the birth rate, death rate, or net diversification (birth minus death) rate of species can be studied [1]. For example, the question, “are nectar spurs a key innovation in plant evolution leading to a rapid radiation?” can be rephrased as, “is the rate of diversification in plant lineages correlated with the presence of nectar spurs?” and this hypothesized association can be tested statistically. In infectious disease phylodynamics, the question “was this intervention effective in containing disease spread?” can be rephrased as, “after the intervention, did the birth rate (effective reproduction number) decrease?” In general, causation is difficult to establish, but the presence of correlation can lend support to hypotheses regarding causes of diversification. Questions involving variation in diversification rates can generally be broken down into two categories. The first class of questions, including the question about nectar spurs, concerns variation in diversification rates across lineages. In these scenarios, models are built that allow the birth and death rates to vary across the branches of the phylogenetic tree [2, 3]. The second class of questions, including the question about intervention efficacy, concerns temporal variation in diversification rates shared by all lineages [4, 5, 6]. In these scenarios, the birth and death rates are modeled as functions of time, but at any instant in time all branches of the tree share a common birth rate and a death rate. This second class of questions and models is our focus in this paper. Our aim is to develop flexible Bayesian nonparametric methods for accurately estimating changes of birth and death rates over time without sacrificing precision.

Birth-death models [7, 8] define a probability distribution on time-calibrated phylogenies— phylogenetic trees where branch lengths are measured in time rather than in evolutionary distances. Early approaches to inferring variability of birth and/or death rates required the use of a time-calibrated phylogeny as data. This involved estimating parameters of birth-death models and then either statistically testing for violations of constant birth and death rates [9] or choosing the best functional form (*e.g.*, two-piece piecewise constant or exponential curves) for birth and death rate trajectories from a set of candidate models via likelihood-ratio tests or the AIC [10, 11]. These early methods had the downside of not accounting explicitly for missing taxa, requiring the use of Monte Carlo simulation in order to determine if the rejection of a constant-rate (or other) model in favor of a more complex model was an artifact of incomplete sampling of phylogenetic lineages [12, 13, 14]. However, the underlying theory and likelihood function for arbitrary functions of birth and death rates including unsampled taxa was already introduced by Nee *et al.* (1994) [8]. Later, the introduction of the piecewise-constant, or episodic, birth-death model (EBD) [15] enabled biologists to perform likelihood-based comparison of birth and death rates’ functional forms while accounting for incomplete taxon sampling (see Höhna (2015) for a review of the EBD and comparison to the work by Nee *et al.* (1994) [16, 8]). The EBD model was extended to work in contexts with serial samples (*e.g.*, fossils) and possibly sampled ancestors [17, 18].

The EBD model divides time into a finite number of intervals and assigns each interval its own set of birth and death rates. The first uses of the EBD model assumed that *a priori* birth and death rates in each interval are independent and identically distributed (iid) [17, 18]. This assumption means that the number of intervals (or epochs) needs to be kept small to keep estimation reasonably precise and to avoid overfitting. Further work on Bayesian modeling using the EBD employed temporally-autocorrelated models derived from discretizing Ornstein-Uhlenbeck and Brownian motion processes [19, 20, 21], which provides smoothing and allows the number of episodic intervals to be larger. May *et al.* (2016) propose another EBD model, where birth and death rates change at an unknown, Poisson distributed number of change-points [22]. Wu (2014) uses a similar change-point model [23]. These random change-point models drastically increase the dimensionality of the parameter space and make it variable, requiring complicated reversible-jump Markov chain Monte Carlo (MCMC) [24] algorithms to sample from the posterior distribution of the number of change-points. However, many other Bayesian nonparametric approaches for estimating functional forms have not been applied to EBD modeling.

Parametric and nonparametric estimation of functional forms is not unique to birth-death processes. For example, population genetics researchers have developed a rich toolbox of Bayesian nonparametric approaches to estimate changes of the effective population size in a neighboring class of coalescent models [25]. In fact, EBD models closely resemble piecewise constant effective population size coalescent models [26, 27, 28]. However, EBD models still lack Bayesian regularization approaches that control the potentially high number of model parameters. For coalescent models, such Bayesian regularization is accomplished by Gaussian Markov random field (GMRF) prior distributions, which underly the skyride [27] and skygrid [28] methods, and by their recently developed analog, the horseshoe Markov random field (HSMRF) [29]. These models provide a rich framework for building more complicated models with covariates [30] and are amenable to computationally efficient MCMC sampling techniques. Our goal is to bring GMRF and HSMRF prior distributions to EBD models and to test their performance.

We implement birth-death models that use GMRF and HSMRF prior distributions for the birth and/or death rates in the statistical phylogenetic software platform `RevBayes` [31]. This implementation allows us to jointly estimate birth-death parameters, phylogeny, and other (nuisance) parameters in a Bayesian framework. We develop an efficient, tuning-parameter-free MCMC algorithm for sampling high dimensional parameter vectors associated with GMRF- and HSMRF-based models. We also devise a framework for setting the global scale parameter—the key parameter controlling the degree of parameter regularization (also called shrinkage)—for both models in terms of the implied prior on the number of “effective” rate shifts. We note that our GMRF-based model is closely related to the work of Duplessis (2016), Condamine *et al.* (2018), and Silvestro *et al.* (2019), who use prior distributions that fall into the class of GMRF distributions, but our work differs from these approaches in important computational and statistical details [19, 20, 21]. Namely, we develop a tuning-parameter free MCMC algorithm that enables efficient exploration of the high dimensional parameter vectors associated with GMRF- and HSMRF-based models and introduce a framework for setting the key hyperprior in an interpretable manner. To the best of our knowledge, this is the first instance of applying HSMRF prior distributions to birth-death processes. We test both GMRF-based and HSMRF-based models on a variety of simulated diversification scenarios, and then apply them to a species-level and an epidemiological dataset. We find that both models are capable of inferring variable diversification rates and correctly rejecting variable models in favor of effectively constant models. In general, in line with previous analyses of HSMRF prior distributions [32, 29], we see that the HSMRF-based model has higher precision than its GMRF counterpart, with little to no loss of accuracy. In empirical applications, we show that these models are useful for detecting a speciation-rate decline in the Australian gecko clade Pygopodidae and a complex pattern of variation in the rate of infection of HIV subtype A in Russia and Ukraine.

## Methods

Our data, **y**, take the form of a multiple sequence alignment. We assume that the alignment **y** has come from the following probabilistic model. First, a tree is generated from a time-varying birth-death process governed by time varying birth rate *λ*(*t*), death rate *μ*(*t*), serial sampling rate *ϕ*(*t*), conditional probability of death upon sampling *r*(*t*) (primarily for phylodynamic applications to represent becoming noninfectious when diagnosed and/or treated), and vector of sampling probabilities **Φ** (with associated sampling times ***t***_**Φ**_, we refer to these as event sampling times). Time starts at 0 at the most recent event (or serial) sampling time and increases into the past, such that the oldest bifurcation in the tree is *t*_*o*_, the time of origin (here also the time of the most recent common ancestor) [8]. We call the resulting reconstructed tree *T*, and it consists only of lineages whose descendants were sampled. For more details on this model, including helpful figures and derivations, see Gavryushkina *et al.* (2014) [18]. On each branch of *T*, evolution proceeds at a rate governed by a molecular clock model [33, 34, 35]. Columns in the sequence alignment evolve independently under a continuous-time Markov chain (CTMC) model, which commonly is referred to as the substitution model. We use the generalized time reversible substitution model [36] with discretized gamma-distributed rate variation across sites (GTR+G) [37]. For notational simplicity we refer to the vector of substitution and clock model parameters as ***θ***, and we discuss the specifics of these models on a case-by-case basis. We can write the phylogenetic likelihood—probability of the alignment under the CTMC substitution model—as Pr(**y** | ***θ***,*T*). All major statistical phylogenetic software platforms can efficiently compute a phylogenetic likelihood via a dynamic programming algorithm, known as the Felsenstein pruning algorithm [38]. We will use the `RevBayes` implementation of this algorithm [31].

In Bayesian inference we need prior distributions for *λ*(*t*), *μ*(*t*), *ϕ*(*t*), *r*(*t*), **Φ**, and *t*_*o*_, as well as prior distributions on ***θ***. We assume that ***t***_**Φ**_ is fixed *a priori* by the user. For our purposes there will only be one time at which event sampling may occur: the present day, making ***t***_**Φ**_ a scalar *t*_Φ_ = 0. Notice that the prior on *T*, conditional on *λ*(*t*), *μ*(*t*), *ϕ*(*t*), **Φ**, and *t*_*o*_, is already specified by the birth-death process. The choice of Pr(*t*_*o*_) depends on the particular group of taxa studied, and the form of Pr(***θ***) on the specifics of the group and the data, so we discuss these on a case-by-case basis. The posterior distribution takes the following form:

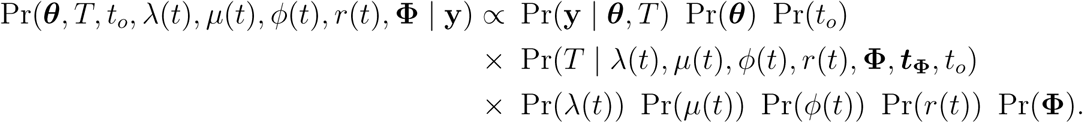

In macroevolutionary analyses including extant species, there is a single event sampling at the present (*t* = 0) with known probability Φ_0_ [39]. In phylodynamic analyses, there may be no event sampling, thus we set Φ_0_ = 0. We make the simplifying assumptions that the serial sampling rate is a constant, *ϕ*(*t*) = *ϕ*, and that the conditional probability of becoming noninfectious upon sampling is a known constant, *r*(*t*) = *r*. For any macroevolutionary dataset *r* = 0, and in our phylodynamic application we assume *r* = 1. Additionally, in our macroevolutionary example there are no serial samples, hence *ϕ* = 0 (in which case *r* is not a parameter of the simplified model). In all analyses we make the additional simplifying assumption that the death rate is a constant *μ*(*t*) = *μ*, and place a mean-0.1 exponential prior on *μ*. In phylodynamic applications, there is often prior information that enables the use of informative prior distributions on *ϕ* and *μ*, which we discuss in a later section. The remaining piece of the puzzle, and our contribution in this paper, is in the specification of Pr(*λ*(*t*)), for which we use Markov random field models. (Note that our implementation and theory of the GMRF and HSMRF can be applied to all time-varying rates; we focused on the birth rate only for simplicity.) Our simplified posterior distribution takes the following form:

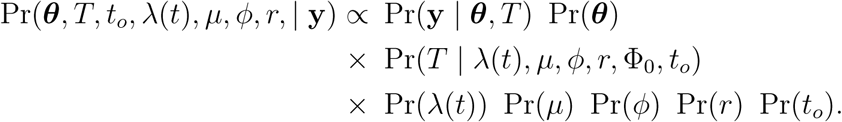

We note that historically Φ_0_ has been called *ρ*, and **Φ** has sometimes been called ***ρ***. However, ***ρ*** has been used to refer to both sampling probabilities [18] and mass extinction probabilities [15, 16, 22], which creates room for confusion.

### Horseshoe Markov random field prior

We define the birth rate on the log scale, *λ*^∗^(*t*) = ln(*λ*(*t*)). Following Stadler (2011), we discretize time into *n* intervals and assume that 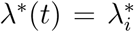 when *t* is in the *i*th time interval, using the parameterization 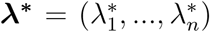 [15]. We find *n* = 100 works well in practice and refer readers to the supplemental materials for a more detailed discussion of the grid size *n*. An HSMRF is a model in which 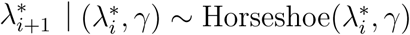, where *γ* is a global scale parameter that controls the smoothness of the overall field. The horseshoe is a distribution used as a shrinkage prior, a statistical tool designed to discern signal from noise [40]. In our case, the HSMRF exerts strong prior belief that 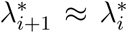; in other words, we do not expect much change in the birth rate between adjacent intervals. However, the horseshoe distribution also has fat (Cauchy-like) tails that allow the HSMRF to behave like a spike-and-slab mixture model [41], giving the HSMRF a property known as local adaptivity. The horseshoe distribution employs an auxiliary variable *σ*_*i*_ and is represented as a scale mixture of normal distributions

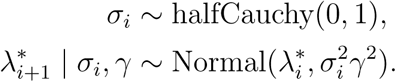

This mixture representation helps explain the local adaptivity of the HSMRF: one or a few (relatively) large changes can be handled by large *σ*_*i*_ without increasing *γ*. We place a Normal(ln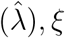) prior distribution on 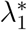, where 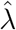 is a rough estimate of net diversification rate. When there are extant lineages in the tree, 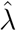 is the maximum likelihood estimator for the net diversification rate, *d*, from Magallon and Sanderson (2001) [42]. When there are no extant lineages in the tree, we put a lower bound on the net diversification rate using the number of births in the tree (excluding the origin or root as appropriate), *B*_*obs*_. The expected net number of births observed in a tree by time *t* is given by 𝔼 (*B*_*t*_) = 2*e*^*t*·*d*^ − 2 if starting at the time of the most recent common ancestor, and 𝔼(*B*_*t*_) = *e*^*t*·*d*^ − 1 if starting with a single lineage. By the method of moments, we can obtain (for the case of starting with the MRCA) 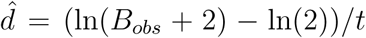, where *t* is the age of the tree. As not all lineages that are born will be sampled in our tree, the number of observed births will be an underestimate of the number of births and our rate will be underestimated, but it will suffice. In practice, when setting the prior for the first birth rate, we use *ξ* = 1.17481, producing, *a priori*, 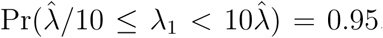. We use a halfCauchy(0, *ζ*) prior on *γ*, where *ζ* is the global shrinkage prior, and we discuss how to set it in a later section. A list of all parameters in the model and their prior distributions can be found in Table 1. The full posterior distribution of our HSMRF-based model parameters is,

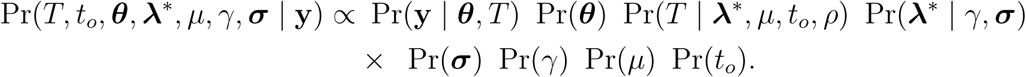

**Table 1.**
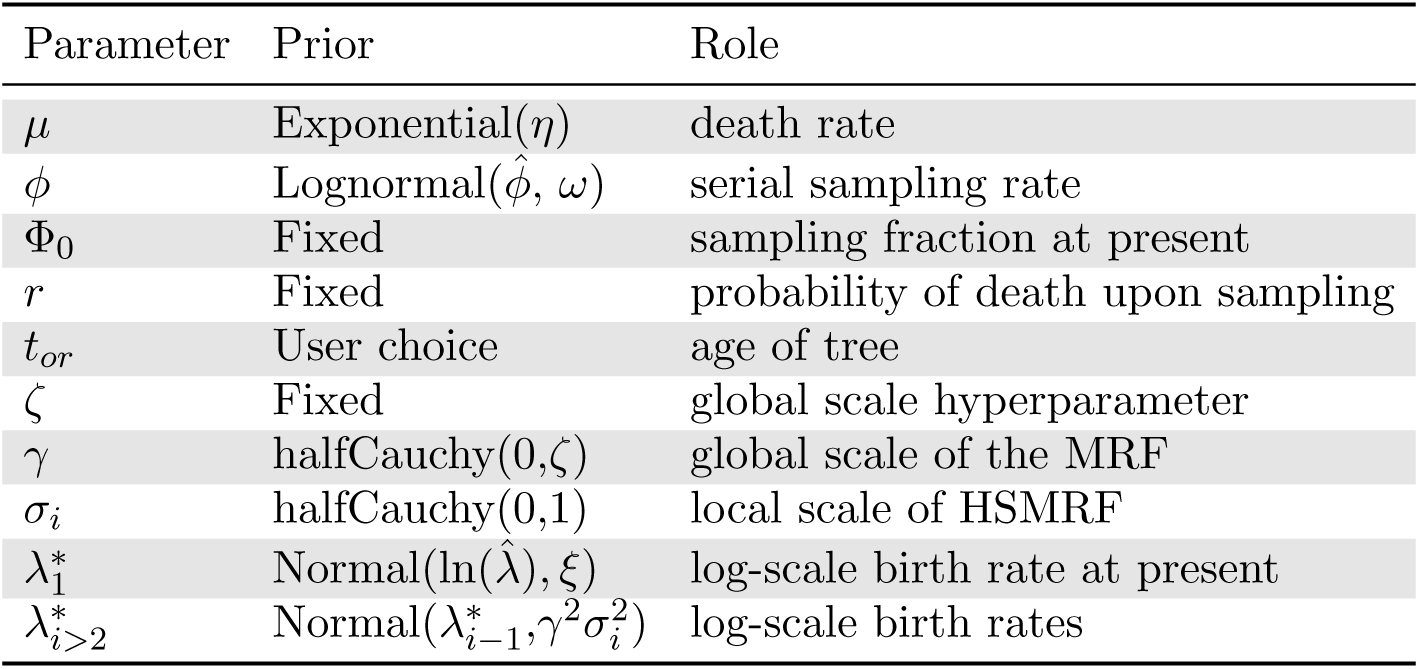
Model parameters, their prior distributions, and their role in the model. In most of our analyses, we assume a constant death rate, *μ*, with an Exponential prior with rate parameter *η* = 10. In phylodynamic applications, there may be more information to set *η*, while in macroevolutionary examples one could instead employ an empirical Bayes approach. When there is serial sampling, we adopt an empirical Bayes approach to setting the prior on the sampling rate, *ϕ*, using a guess at the tree age and the number of tips to obtain 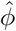. In practice we set *ω* = 1.17481. In analyses without serial sampling, *ϕ* = 0. For details on computing 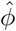, see the supplemental materials. The sampling fraction at present, Φ_0_ and the probability of death upon sampling, *r* are taken to be known *a priori*. The age of the tree, *t*_*or*_ is fixed to the observed height if the tree is data, else it is a variable with the prior determined by the user. For models with *n* = 100 intervals, we set *ζ* = 0.0021 for HSMRF-based models and *ζ* = 0.0094 for GMRF-based models, while for models with other *n*, we provide code for setting *ζ*. The GMRF-based model lacks local scale parameters ***σ***. We adopt an empirical Bayes approach to setting the prior on the first log-birth-rate using a guess at the tree age and the number of tips to obtain 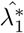. In practice we set *ξ* = 1.17481. In models where the death rate varies, the previously discussed prior on *μ* serves as the prior on *μ*_1_, and the rest of the prior is accomplished via an MRF model exactly as with the birth rate.

| Parameter | Prior | Role |
| --- | --- | --- |
| $\mu$ | Exponential($\eta$) | death rate |
| $\phi$ | Lognormal($\hat{\phi}, \omega$) | serial sampling rate |
| $\Phi_0$ | Fixed | sampling fraction at present |
| $r$ | Fixed | probability of death upon sampling |
| $t_{or}$ | User choice | age of tree |
| $\zeta$ | Fixed | global scale hyperparameter |
| $\gamma$ | halfCauchy(0, $\zeta$) | global scale of the MRF |
| $\sigma_i$ | halfCauchy(0, 1) | local scale of HSMRF |
| $\lambda_1^*$ | Normal($\ln(\hat{\lambda}), \xi$) | log-scale birth rate at present |
| $\lambda_{i>2}^*$ | Normal($\lambda_{i-1}^*, \gamma^2 \sigma_i^2$) | log-scale birth rates |

We approximate the above posterior distribution using the following MCMC strategy. We use standard Metropolis-Hastings kernels available in `RevBayes` to update the tree *T*, time of the root *t*_*o*_, substitution model parameters ***θ***, and extinction rate *μ*, and the first log-speciation rate, 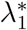 (see Höhna *et al.* (2017) for a description of the standard `RevBayes` Metropolis-Hastings kernels [43]). Since vectors ***λ***^∗^ and ***σ*** can be high dimensional, we update the vectors in blocks. First, we reparameterize the model to work with the first order differences **Δ** instead of ***λ***^∗^ (see Figure 1). This allows us to sample the vector **Δ**, where all elements are *a priori* independent, instead of the vector ***λ*** where the adjacent values are highly correlated, greatly increasing the efficiency of the MCMC. Further, under the GMRF and the hierarchical representation of the HSMRF, all the **Δ** are Normal random variables, enabling us to employ an elliptical slice sampler [44] for **Δ** = (Δ_1_,*…*, Δ_*n*−1_). The (conditional) normality of the **Δ** also allows us to employ a Gibbs sampler for *γ* and ***σ***, which allows us to adequately sample the tails of the posterior distribution. Without this elliptical slice sampler and Gibbs sampler combination, MCMC for these models fails to converge to the posterior distribution. We defer a more thorough discussion of our MCMC strategy to the supplemental methods. We also note that there are directionless specifications of MRF models which make it implicit that information is shared across adjacent intervals in both directions (that is that the full conditional distribution of 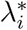 depends on both 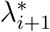 and 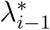 For more on this subject, we direct readers to Rue and Held (2005) [45].

**Figure 1.**
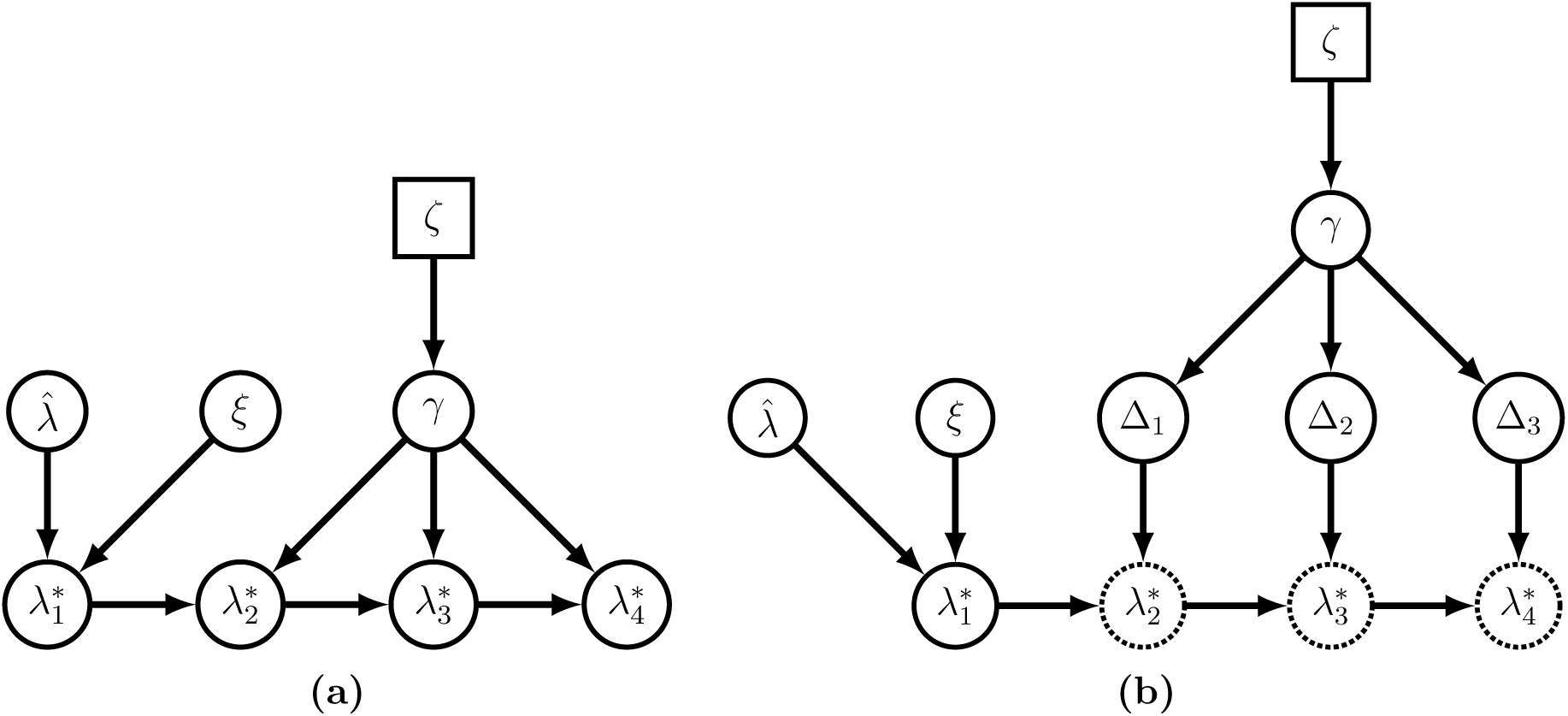
Simplified versions of our MRF-based models, shown as a grid of size 4. To highlight the structural similarities between the GMRF- and HSMRF-based models, we draw the directed acyclic graph (DAG) as if we had an analytical form of the horseshoe distribution (that is, we omit the local scale parameters of the HSMRF). In (a), we show the idealized general MRF model, while in (b), we show how we can reparameterize the model in terms of a vector **Δ** of independent random variables. From **Δ**, we can recover ***λ***^∗^ as 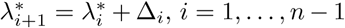. This reparameterization greatly improves the efficiency of MCMC sampling. When drawing the model as a DAG, squares represent constant values, closed circles stochastic values, and open circles deterministic transformations of other nodes.

### Gaussian Markov random field prior

Our GMRF-based model can be seen as a special case of an HSMRF-based model where *σ*_1_ = ··· = *σ*_*n*_ = 1, meaning 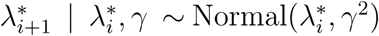. The lack of local scale parameters makes the GMRF-based model a non-locally-adaptive model. For the GMRF-based model, the posterior distribution is

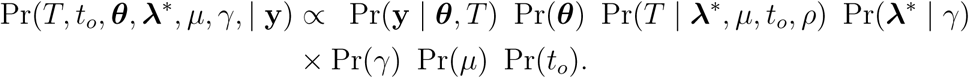

The MCMC algorithm to approximate the above posterior distribution is the same as for the HSMRF-based model, except we do not need to update the vector ***σ***.

### Setting the prior on the global scale parameter

In both HSMRF- and GMRF-based models, the global scale parameter, *γ*, controls the smoothness of the overall field, with smaller values favoring less variability. Following Faulkner *et al.*, we take a hierarchical approach and place a prior distribution on the global scale parameter, such that *γ* ∼ halfCauchy(0, *ζ*) [29]. We choose *ζ* in terms of *s*_*e*_ — the number of “effective shifts” in the birth rate and we define an effective shift to be an event {*λ*_*i*+1_*/λ*_*i*_ *<* 1*/ϵ* or *λ*_*i*+1_*/λ*_*i*_ ≥ *ϵ*}. That is, an effective shift is the event where two adjacent birth rates are different by more than -fold. We set *ϵ* = 2, reflecting that a 2-fold change in the birth rate is biologically meaningful and statistically detectable. Setting *ζ* is then accomplished implicitly by setting the prior expected number of effective shifts, 𝔼 [*s*_*e*_], which is more interpretable than *ζ*. In this setup, 𝔼 [*s*_*e*_] is the expectation of a binomial random variable with probability *p* that there is an effective shift between *λ*_*i*+1_ and *λ*_*i*_. Since we can compute *p* given a particular value of *ζ*, and since there is no obvious closed form solution, we use numerical methods to solve for *ζ*. Code to calculate *ζ* from 𝔼 [*s*_*e*_] is available in the GitHub repository, see section “Code and data availability.”

We find that in practice 𝔼 [*s*_*e*_] = ln(2) produces a prior that is reasonably conservative yet flexible. This yields *ζ*_*HSMRF*_ ≈ 0.0021 for HSMRF-based models and *ζ*_*GMRF*_ ≈ 0.0094 for GMRF-based models. An alternative approach to specify *ζ* examines the implied prior distribution on *λ*_*n*_*/λ*_1_, *i.e.*, the prior distribution on the fold change across the entire process. *A priori*, for the HSMRF on a grid size *n* = 100, 𝔼 [*s*_*e*_] = ln(2) leads to Pr(0.5 ≤ *λ*_*n*_*/λ*_1_ *<* 2) ≈ 0.76 and Pr(0.1 ≤ *λ*_*n*_*/λ*_1_ *<* 10) ≈ 0.9. While we do not use this approach to set *ζ*, it shows that our chosen value for *ζ* focuses the prior mass on reasonable regimes while leaving room for rather substantial amounts of change. For completion, in the supplementary materials we provide more context for these choices of prior, including an alternative choice of 𝔼 [*s*_*e*_] following Drummond and Suchard (2010), and examine two additional frameworks, bounding the marginal variance of the GMRF and HSMRF (explored by Sørbye and Rue (2014) and Faulkner *et al.* (2018)), and bounding the effective number of parameters in the model (explored by Piironen and Vehtari (2017)) [46, 47, 29, 48].

## Results

### Simulation study

To understand statistical properties of both random field birth-death models, we perform a (nonexhaustive) simulation study. Some of the most debated questions in species diversification concern diversification-rate decreases [49, 50, 9, 51, 52, 53, 54], and the ability to detect effective epidemiological interventions hinges on the ability to accurately estimate decreases in the rate of infection, so we consider simulation scenarios where the birth-rate declines through time. We devise a series of piecewise-linear functions *λ*(*t*) in which the birth rate decreases through time. For each model, we use the R package `TESS` [55] to simulate 100 trees conditioned on the tree age (*t*_*o*_ = 100), with complete species sampling (Φ_0_ = 1), and choosing values for *λ*(*t*) and *μ* to give an expectation of 200 species/tips at the present. Given the underdeveloped infrastructure for simulating serially-sampled trees, we focus on trees where all samples are at the present day (*ϕ* = 0), but see Barido-Sottani *et al.* (2019) for recent developments [56]. When analyzing these simulations, we take the tree and sampling fraction to be known. Treating the tree as data allows us to focus on the performance of the random field birth-death models without worrying about potential sources of bias during time-calibrated tree estimation [57]. Taking the tree as data also mirrors the predominant historical usage of models of rate variation, detecting variation in trees previously inferred [11, 2, 3, 15].

We assess model performance by looking at four summaries of the inferred birth-rate trajectories. We take as our estimate of *λ*(*t*) the birth-rate trajectory defined by the median posterior of each birth rate *λ*_*i*_. First, to quantify bias we use the Mean Absolute Deviation (MAD) of the estimated birth-rate trajectory, given by 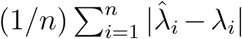. Second, we look at the Mean Absolute Sequential Variation (MASV) of the estimated trajectory, the gross change inferred, given by 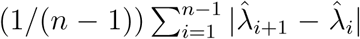. Where the simulated trajectory is variable, it is more useful to consider the relative MASV (RMASV), MASV(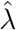)/MASV(*λ*). If RMASV *>* 1, the inferred trajectory is more variable than the true trajectory, and if RMASV *<* 1, it is less variable. Third, we look at the fold change (FC) of the estimated trajectory, 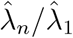. This will show us if we capture the presence of an overall change in the birth rate, even if we fail to capture the specific pattern. Finally, we look at the average width (across all estimated birth rates, *λ*_*i*_) of the 90% posterior credible interval as a measure of precision, 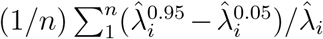. This measure, which we call relative precision (RP), is both more interpretable than the raw credible interval and more comparable across simulations.

#### Constant-rate simulations

Our first simulations are from a constant-rate diversification model, such that *λ*(*t*) = *λ*. This allows us to test the tendency towards what could be termed “false positives,” the detection of spurious rate variation. Both the GMRF and HSMRF birth-death models can produce effectively constant-rate trajectories, though their flexibility is not without minor drawbacks. Ridge plots of performance measure histograms across all simulations are shown in Figure 3. The trajectories estimated by both models have low MAD performance measure, FC ≈ 1, and small RP performance measure, indicating generally good performance. Further, compared to fitting constant-rate models, the increase in the MAD performance measure from inferring the variable-rate models is negligible, and the decrease in precision is small (Supplementary Figure S1). Thus, the primary drawback to using these models to fit constant-rate trajectories is that there are false positives. In other words, the models occasionally fit trajectories where the inferred change between the beginning and end of the process does not appear negligible. However, comparisons to the prior make it quite clear that both the GMRF and HSMRF are fitting effectively constant-rate trajectories. For both models roughly 99% of the prior MASV is greater than 0.01, while roughly 5% of the posterior MASV is greater than 0.01, further indicating that the models are producing effectively constant trajectories. Computing such a significance threshold to reject a constant rate model can be computed using Monte Carlo simulation [58]. The HSMRF and GMRF produce very similar average error and fold change, though the distributions of these metrics for the GMRF are more tightly focused around the target values (MAD of 0, FC of 1), and the GMRF has slightly tighter credible intervals. The GMRF generally estimates narrower credible intervals, while the HSMRF generally estimates trajectories with lower MASV.

**Figure 2.**
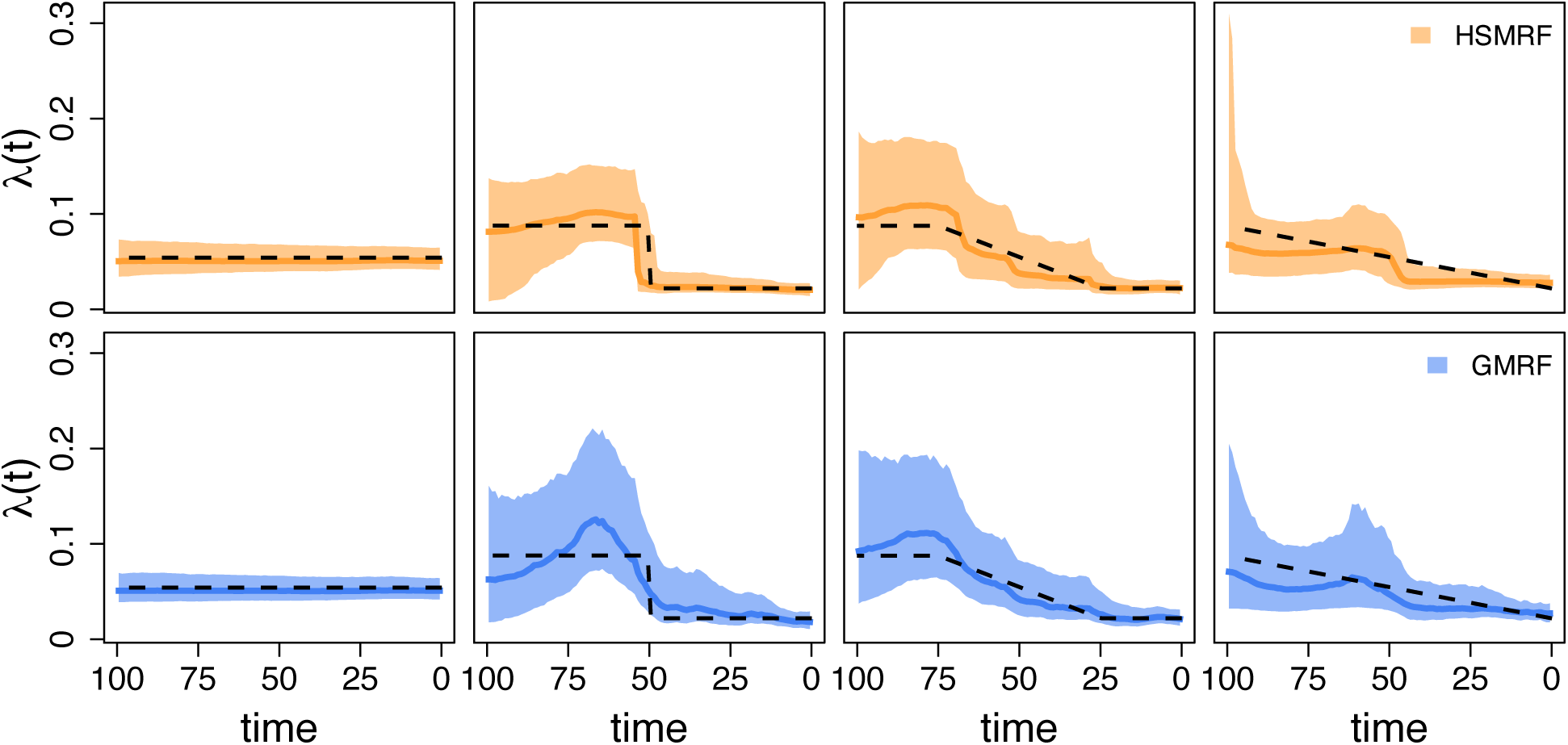
Inferred birth-rate trajectories from four individual simulations. The dashed line is the true simulating birth rate, the dark colored line is the posterior median trajectory (the median is taken separately for each grid cell), and the shaded region show the 90% Credible Interval (CI) for the rate. The leftmost column is from the constant-rate simulations, and the right three columns demonstrate the effect of changing the shift duration (the length of the tree over which the birth rate changes), from an instantaneous shift to a constant change model. When we focus instead on the location of the shift, all simulations are piecewise-constant as in the second column. In each column, we show the simulation with the most average performance measured in terms of the Mean Absolute Deviation of both the GMRF and HSMRF.

**Figure 3.**
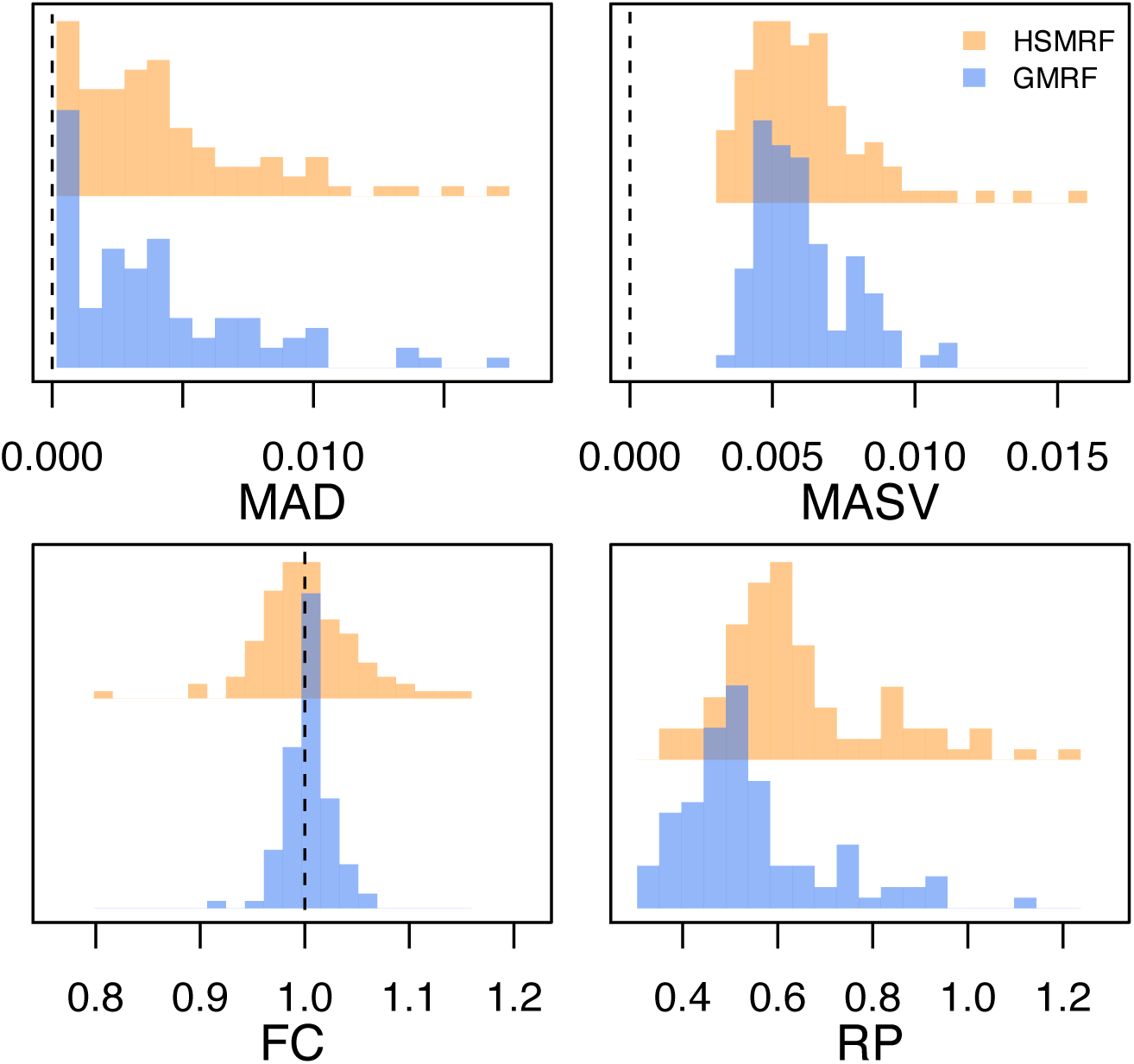
Performance of the models on simulated constant-rate datasets. MAD (Mean Absolute Deviation) measures the error in the estimated trajectory. MASV (Mean Absolute Sequential Variation) measures the total amount of change relative in the trajectory, horizontal line at true value for reference. FC (Fold Change) measures the fold change from present to past, dashed line at true value for reference. RP (Relative Precision) is a measure of precision, the average width of the 90% Credible Interval relative to the birth rate.

#### Piecewise linear simulations

Our primary time-variable simulations examine the impact of the shift duration, *i.e.*, the amount of time over which the birth-rate changes. To examine this, we build a piecewise-linear birth-rate function, where the birth rate is *λ*_1_ for 100 ≥ *t > t*_1_, *λ*_2_ for *t* ≤ *t*_2_, and a linear interpolation for *t*_1_ ≥ *t > t*_2_. We center the shift at 50 ((*t*_1_ + *t*_2_)/2 = 50), and simulate shift durations (*t*_2_ − *t*_1_) of 0%, 25%, 50%, 75%, and 100% of the age of the tree. All simulation parameters are recorded in Supplementary Table S1. The HSMRF-based model performs better when the shift is fast (when *t*_2_ − *t*_1_ is small), and the GMRF-based model performs better when the shift is slow (when *t*_2_ − *t*_1_ is large, Figure 4a). For the HSMRF-based model, the MAD performance measure is lower and both the MASV and FC performance measures are closer to the truth when the true shift is shorter. In contrast, for the GMRF-based model, the MASV performance measure gets closer to the truth, *i.e.*, the error decreases, and the RP performance measure gets narrower as the shift duration increases. In some simulations, both models effectively fit constant-rate trajectories. The HSMRF-based model also has a tendency towards fitting steep shifts even in cases where the true shift is slow (Figure 2). Both models though have difficulty with continuous, slow declines where they have a tendency to underestimate the total change. However, comparisons to the prior show that both models tend to detect some change. For both the HSMRF- and GMRF-based models, the median FC is approximately 2 for the continuous decline simulations (52% and 54% of FC are greater than 2, respectively). Simulations from the prior of the HSMRF-based model show that 12% of trajectories are expected to have a larger fold change than 2, while for the GMRF-based model 3% are expected to be greater than 2. Thus, compared to the prior, the posterior median trajectories have shifted to larger fold changes. Further, while the prior median fold change is 1, only 3% and 2% of FC inferred by the HSMRF- and GMRF-based models are below 1. All of this indicates evidence for rate variation.

**Figure 4.**
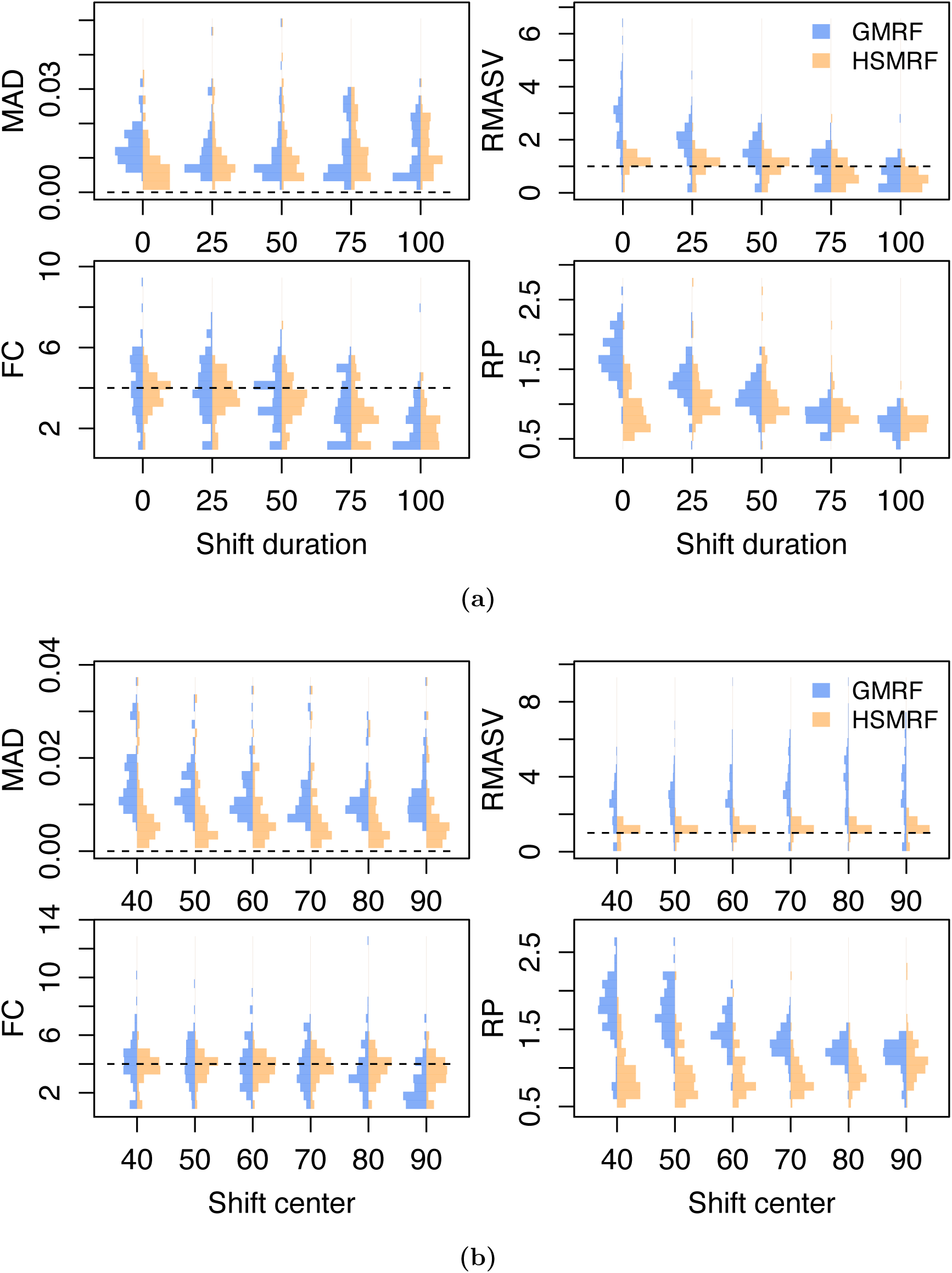
The effect on parameter inference of (a) changing the (four-fold) rate shift from instantaneous to the entire length of the trajectory and (b) changing the center of an instantaneous (four-fold) rate shift. MAD measures the error in the estimated trajectory. RMASV (Relative MASV) measures the total amount of change relative to the true MASV, horizontal line at 1 for reference. FC measures the fold change from present to past, dotted line at true value for reference. RP is a measure of precision, the average width of the 90% Credible Interval relative to the birth rate.

#### Piecewise constant simulation

We also examine the effect of varying the location of the instantaneous birth-rate shift. To do this, we build a piecewise-constant birth-rate function, where the birth rate is *λ*_1_ for 100 ≤ *t < t*_*shift*_, *λ*_2_ for *t* ≤ *t*_*shift*_; we simulate *t*_*shift*_ = 90, 80, *…*, 40 (*e.g.*, Figure 2 second panel). The location of the rate shift should influence the capacity for detection by altering the expected number of births in the pre-shift portion of the tree. As the shift moves from past to present, for the HSMRF-based model the MAD performance measure decreases and the RP performance measure gets smaller (Figure 4b). However, for the GMRF-based model, as the shift becomes more recent, the RMASV performance measure becomes increasingly inflated, indicating trajectories that are too variable. This is due to the GMRF-based model estimating rather substantial variation in the more ancient portions of the tree. The HSMRF-based model outperforms the GMRF-based model in most performance measures and for most shift locations.

#### Shift magnitude

The magnitude of the birth-rate shift should also impact the capacity for detection, so we simulate shifts of two magnitudes for all scenarios outlined above. For our low magnitude shift, we simulate a two-fold change, and for our larger shift, we simulate a four-fold change. Unsurprisingly, it is harder to detect smaller shifts. Results for different functional forms are qualitatively similar between shift magnitudes, so we present only the results for the four-fold shifts in Figure 4. In many cases with two-fold shifts, the inferred trajectory is effectively constant. Thus in general the MAD performance measure is higher, the RMASV, FC, and RP performance measures are lower. Supplementary Figures S4 and S5 give full simulation results for the two-fold case comparable to Figure 4.

### Time-varying death rates

We also investigated the ability of our models to simultaneously infer both time-varying birth and death rates. We devised a piecewise-constant simulation scenario loosely based on a dataset of Hepatitis C infections in Egypt, which is often used as a benchmark for assessing phylodynamic methods [17, 29]. In this scenario, the death rate experiences an approximately five-fold increase, while the birth rate first undergoes a five-fold increase and then a ten-fold decrease. Changes in birth and death rates are asynchronous. We simulate 100 trees with isochronous sampling (Φ_0_ = 1), targeting an expectation of 200 tips as in the main simulation study. We additionally simulate 100 trees with heterochronous sampling (*ϕ* = 0.009,*r* = 1) using `TreePar` [15], producing trees with an average of 175 tips. For comparability (in terms of the expected number of extant taxa at any time in the tree), in the heterochronous simulations we adjust *μ*(*t*) such that the total death rate (*μ*(*t*)+ *ϕr*) is the same as the death rate (*μ*(*t*)) in the isochronous simulations.

We analyze both tree sets with our GMRF-based and HSMRF-based models in two different analysis setups. First, we attempt to infer both *λ*(*t*) and *μ*(*t*). Second, we misspecify the true model by inferring *λ*(*t*) while inferring a constant *μ*. In Figure 5 we show representative estimates of birth and death rates for both models and both analysis setups. We also summarize our results using the MAD, RP, and RMASV performance measures in Figure 6. We do not report fold change here as it is not a useful measure here. This is because the birth-rate trajectory has more than one shift, and in some intervals *μ*(*t*) = 0, making the fold change infinite (this choice of *μ*(*t*) allows us to keep the total death rate the same between the heterochronously sampled and isochronously sampled simulations).

**Figure 5.**
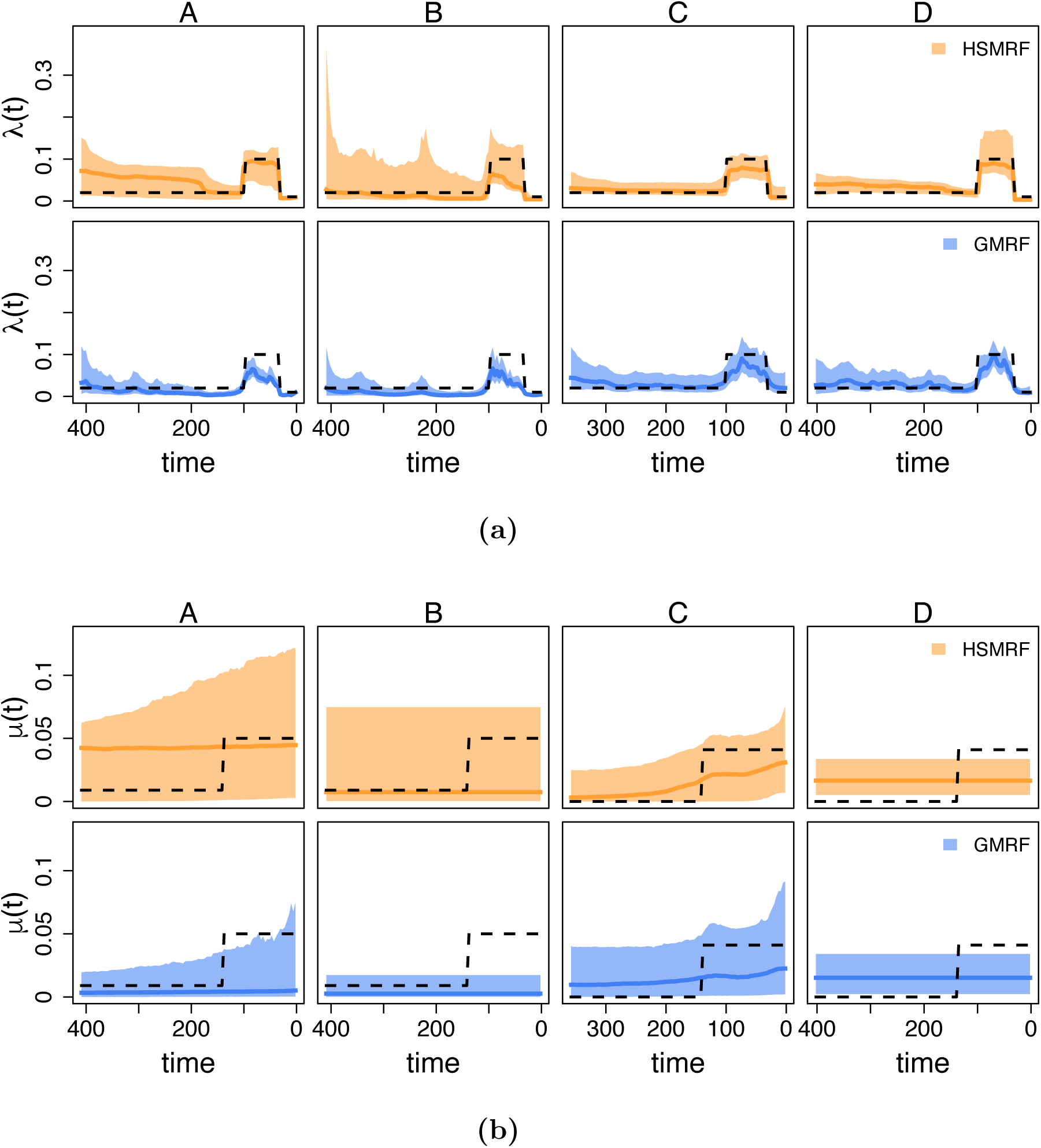
Inferred (a) birth-rate trajectories and (b) death-rate trajectories from four individual simulations with time-varying birth and death rates. The dashed line is the true simulating rate, the dark colored line is the posterior median trajectory (the median is taken separately for each grid cell), and the shaded region show the 90% Credible Intervals (CIs) for the rate. In each column, we show the simulation with the most average performance measured in terms of the Mean Absolute Deviation of the birth- and death-rate trajectories from both the GMRF and HSMRF (columns are shared across birth-rate and death-rate subfigures). The column labels A, B, C, and D identify the different combinations of tree simulations and analysis setup. A and B are analyses of trees with isochronous sampling, C and D heterochronous sampling. A and C are analyses where time-varying death rate, *μ*(*t*) is inferred, B and D where a constant death rate, *μ*, is inferred.

**Figure 6.**
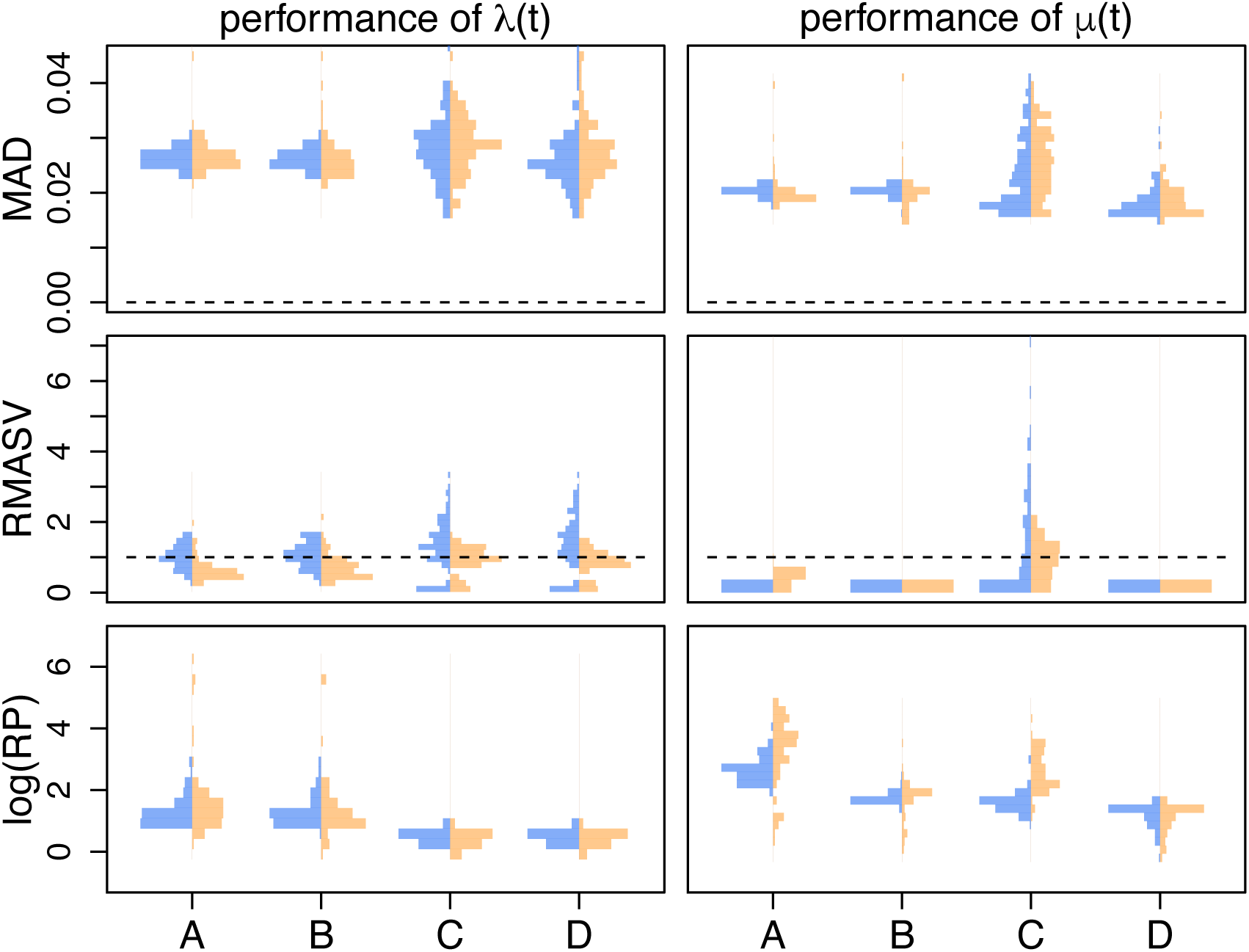
Performance of the models on simulated datasets where both the birth- and death-rate trajectories. MAD measures the error in the estimated rate. RP is a measure of precision, the width of the 90% Credible Interval relative to the true rate. RMASV measures the total amount of change relative to the true MASV, horizontal line at 1 for reference. The column labels A, B, C, and D identify the different combinations of tree simulations and analysis setup. A and B are analyses of trees with isochronous sampling, C and D heterochronous sampling. A and C are analyses where time-varying death rate, *μ*(*t*) is inferred, B and D where a constant death rate, *μ*, is inferred.

First, we examine the ability of our models to infer time-varying death rates. Our simulations reveal that estimating time-varying death rates can be quite difficult, especially without serial samples. Without heterochronous sampling, there is very little signal in any analysis for time-varying death rates; the performance is essentially equivalent to fitting a constant death rate. When heterochronous samples are present, it is possible to detect time-varying death rates and estimates become more accurate. However, even with heterochronous samples both models frequently underestimate the variability of the death-rate trajectory as measured by the RMASV performance measure. The RMASV performance measure also shows that the HSMRF-based model is much better at detecting appropriate amounts of variation than the GMRF-based model.

We additionally investigate the ability of our models to infer time-varying birth rates in the presence of a varying death rate, and find that estimating time-varying birth rates is more difficult when the death rate varies. The MAD performance measure is higher in the time-varying death simulations than in the main simulation study and the RP performance measure is larger. In the absence of serial samples, the RMASV performance measure shows that the amount of variability is generally underestimated, though there is still clear evidence for variation. As the trajectories have similar birth rates to the constant-death simulations, the MAD performance measures should be largely comparable, despite not being relative measures like the RP and RMASV performance measures. The presence of serial samples greatly improves estimation of the total variability of the trajectory as measured by the RMASV performance measure, and greatly improves accuracy and precision. Fitting a model with a constant death rate generally has little effect on how well the birth rate trajectory is estimated.

### Empirical analysis of Pygopodidae

Pygopodidae is a clade of approximately 46 legless geckos [6]. Recently, Brennan *et al.* (2017) used several birth-death models to investigate the history of diversification in this group, examining trends in speciation over time using a posterior sample of 100 phylogenies estimated via BEAST 1.8.3 [6, 59]. The majority of their analyses revealed a drastic speciation-rate decrease in the recent (2-5 million years) past, though there was some disagreement between methods over the significance and timing of the shift. Here we revisit the question of significance and timing of the birth-rate shift in full joint analyses of phylogeny and both our GMRF and HSMRF birth-death models from molecular sequence data. In these analyses, we assume a constant death rate, *μ*. Details of the substitution and clock models are available in the supplemental materials, as are details of MCMC convergence diagnostics performed.

Our dataset includes 41 out of 46 representatives of Pygopodidae, which we use to set the species sampling fraction, Φ_0_ = 0.89. We employ calibrations on the same nodes as Brennan *et al.* (2017), resulting in a calibration for the root node and one each on the genera *Delma* and *Apprasia* [6]. Following Brennan *et al.* (2017), we place a Uniform(19.5, 29.0) prior on the root age [6]. To set up our grid, we thus choose to divide the interval [0, 29] into 100 intervals/epochs of equal length.

GMRF- and HMRF-based models produce a clear visual signature of a diversification-rate decrease (Figure 7), with a higher rate from the origin of the clade up until at least 12 Ma, and a lower rate afterwards. The HSMRF-based model favors a steeper decrease ending approximately 6 Ma, while the GMRF-based model favors a much slower decline that starts approximately 14 Ma and lasts until approximately 2 Ma. Over the range [2 Ma, 12 Ma], the HSMRF estimates a 3.43-fold decrease (90% Credible Interval (CI) [1.12, 8.49]), while the GMRF estimates a 2.41-fold decrease (90% CI [1.00, 7.58]). The HSMRF produces 90% credible intervals for the speciation rate that are generally narrower than the GMRF-based model’s intervals. The behavior of both models is in line with the simulation results for fast to intermediate shifts, with the HSMRF inferring a faster shift of larger magnitude with tighter credible intervals than the GMRF-based model.

**Figure 7.**
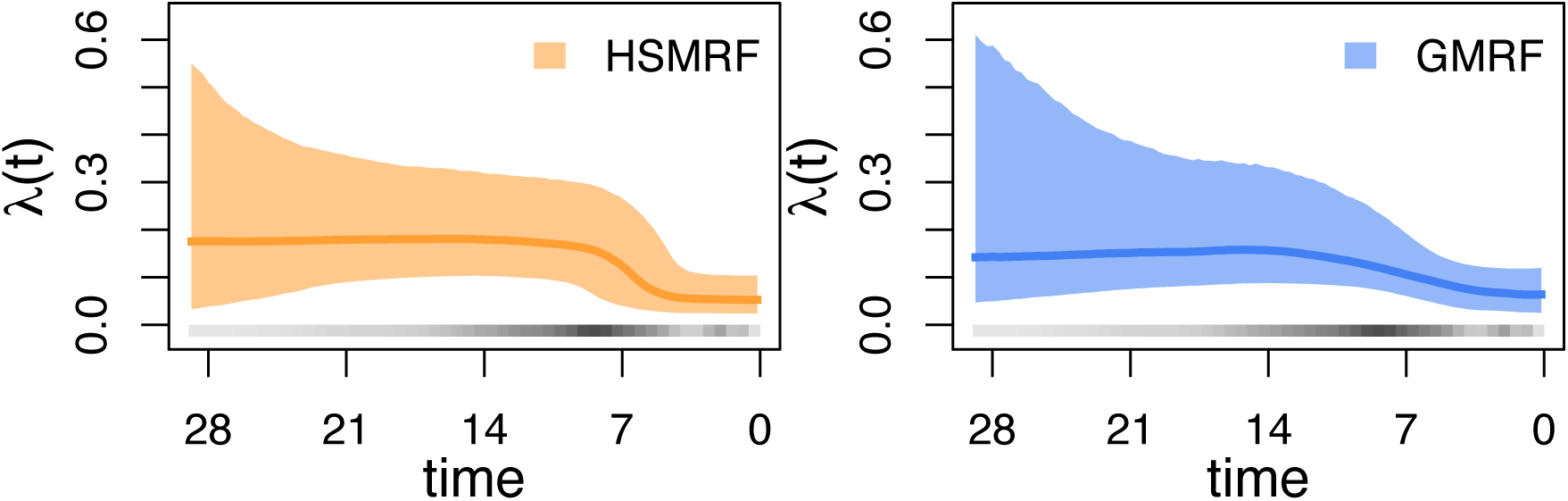
Analyses of the Pygopodidae dataset. Plotted are posterior median trajectories (dark lines) and 90% credible intervals (shaded regions). Time is in millions of years before the present day. In grey is a heatmap of the inferred divergence times.

Given that the posterior distributions of adjacent birth rates are highly correlated, testing for a shift in a specific interval from *λ*_*i*_ to *λ*_*i*+1_ could suggest there is no shift even when there is clearly a shift present in the overall trajectory. However, we can avoid this issue by testing hypotheses over longer timespans. The Bayes factor [60] in support of an *s*-fold decrease between *t*_*start*_ and *t*_*end*_ is given by,

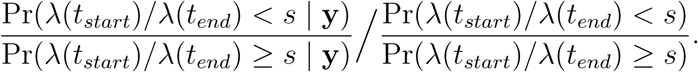

For an *s*-fold increase, the inequalities are reversed. If we are interested in the evidence of a shift over the range [2 Ma, 12 Ma], we can compare the speciation rates in the appropriate intervals for a given shift size *s*. For our grid, the 7th interval ends at 2.03 Ma, while the 43rd starts at 12.18 Ma, and we would test hypotheses regarding *λ*_7_*/λ*_43_. Then all we need to know are the posterior and prior probabilities of observing a shift of at least *s* (or less than *s* if testing a decrease). If we were instead interested in testing simply for the presence of a shift, then we choose *s* = 1. Under both our HSMRF- and GMRF-based models, the prior probability Pr(*λ*_*i*_*/λ*_*j*_ *<* 1 | *HSMRF*) = 0.5 (for any *i* ≠ *j*), making the denominator (the prior odds) 1 and only requiring us to compute the numerator (the posterior odds). For the HSMRF-based model, Pr(*λ*_7_*/λ*_43_ *<* 1.0 | **y**, *HSMRF*) = 0.98, and the 2ln(BF) in favor of a birth-rate shift over this interval is 7.73 (using the nomenclature of Kass and Raftery (1995), “strong” support [60]). For the GMRF-based model, equivalent calculations produce a 2ln(BF) in favor of a birth-rate shift of 5.53 (“positive” support). If we had instead been interested in testing for a shift of a particular magnitude, we could simulate under the prior to estimate the prior odds.

### HIV Dynamics in Russia and Ukraine

In Eastern Europe and Asia, the use of injected drugs was a driving force in HIV epidemics for many years and continues to be an important factor in the spread of HIV [61]. Russia and Ukraine have a particularly high number of people who inject drugs, 2 million individuals combined, and a total of 1 million HIV-infected individuals [62]. These factors, plus a limited effort to reduce the scope of the problem in the beginning of the epidemic, make Russia and Ukraine a good source of data for estimating how HIV spreads among those who inject drugs. Vasylyeva *et al.* (2016) used a number of approaches, including phylodynamic methods, to study the course of the epidemic from the 1980s through 2011 [62]. They estimated that half of all secondary infections take place during the first month post-infection. They further identified a massive increase in the size of the infected population during the 1990s, and estimated that during this period each newly infected individual transmitted to at least 5 others.

When using birth-death models for infectious disease phylodynamics, the primary parameter of interest is the effective reproductive number at time *t, R*_*e*_(*t*). This is defined as the average number of individuals who will be infected by a single infectious individual introduced into a population with the same numbers of susceptible and removed individuals as are present in the population of interest at time *t* [63]. In a constant-rate birth-death-sampling-treatment process, the average duration of an infection is the inverse of the total rate of becoming noninfectious, or (*μ* + *rϕ*)^−1^. The expected number of infections an individual will cause over a timespan *t* is given by *λ* · *t*, approximately for small *t*. Thus, in the constant-rate case, the expected number of secondary infections caused by an individual is *R*_*e*_ = *λ/*(*μ*+*rϕ*). In the time-varying case, if an individual becomes infected at time *t*, we use the rates at that time to compute the expectation and obtain *R*_*e*_(*t*) = *λ*(*t*)/(*μ*(*t*)+ *r*(*t*)*ϕ*(*t*)).

To understand the dynamics of HIV in Russia and Ukraine in the time period of interest, we use the sequence alignment for the *env* region from Vasylyeva *et al.* (2016) [62]. We analyze this dataset (457 sites for 92 sequences) under both the HSMRF-based and GMRF-based models, defining 2011 (the time of the most recent sample) to be the present day and dividing the range [0,29.1] into 100 evenly-sized intervals. We employ a Normal(29.1,5.0) root age prior, truncated to be older than the oldest sample age of 18. We fix *r* = 1, corresponding to the assumption that an individual, once sequenced and diagnosed, will not cause any further infections because they will be provided treatment and will have undetectable viral load. As there is information about the duration of infection in HIV, and thus the death rate, we replace our usual Exponential(10) prior with a Lognormal(−2.272,0.073) prior on the death rate, *μ* (the rate of becoming noninfectious in the absence of sampling and treatment). This corresponds to an *a priori* 95% probability that an untreated individual will be infectious for between 8.4 and 11.2 years [62, 64]. Details of the substitution and clock models are available in the supplemental materials, as are details of MCMC convergence diagnostics performed.

While the model we fit only has a time-varying birth rate, we plot the more informative *R*_*e*_(*t*) instead in Figure 8. Both the HSMRF-based and GMRF-based models show evidence for a spike in *R*_*e*_(*t*) in the early 1990s and a sharp decrease at the end of the 1990s. We quantify support for shifts in *λ*(*t*) instead of *R*_*e*_(*t*), as we do not directly parameterize the effective reproductive number. The 2ln(BF) in favor of an increase between 1992 and mid-1994 (of any magnitude) are 6.20 (strong support) for the HSMRF-based model and 8.06 (strong support) for the GMRF-based model. Similarly, the 2ln(BF) in favor of a decrease between 1999 and 2001 are 7.63 (strong support) for the HSMRF-based model and 9.91 (strong support) for the GRMF-based model. However, where the HSMRF-based model largely shows evidence for a consistently elevated rate in this period, the GMRF-based model shows a sharp dip midway through the decade, with the 90% CI including *R*_*e*_(*t*) = 1. The HSMRF-based model estimates an average rate in this interval of 3.99, with rates that may be as low as 1.83 or as high as 7.88, and the GMRF-based model estimates an average rate of 3.61 with rates possibly as low as 0.55 or as high as 11.65.

**Figure 8.**
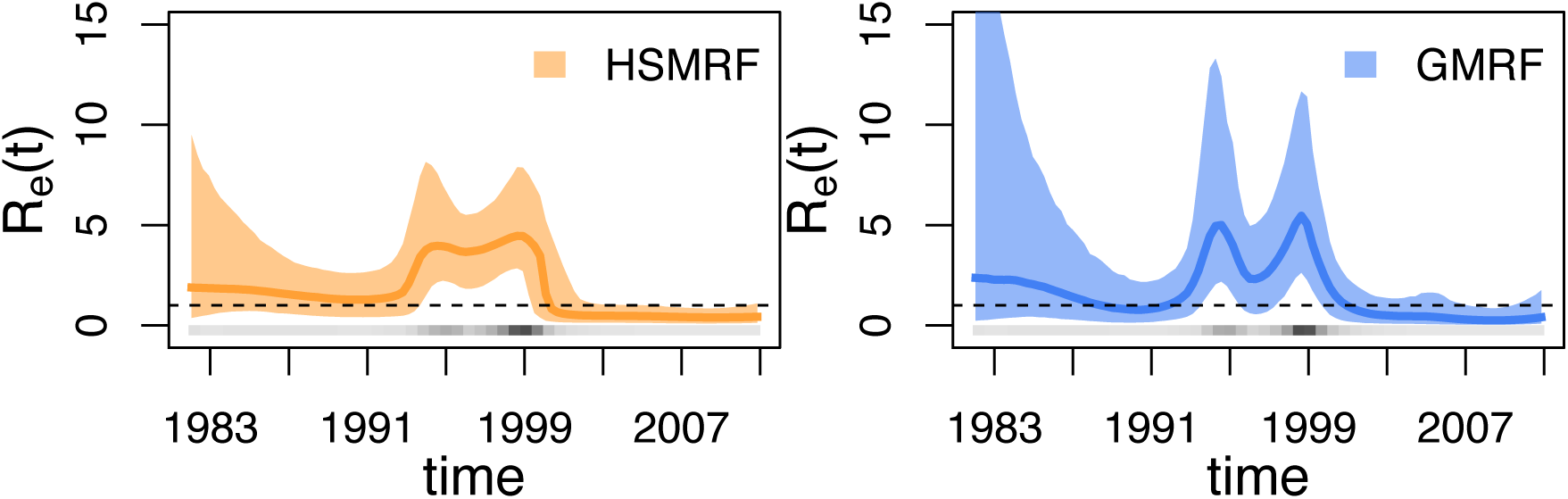
Analyses of the HIV dataset. Plotted are posterior median trajectories (dark lines) and 90% credible intervals (shaded regions). The upper CI for the GMRF-based analysis extends to ≈26, we have truncated the figure for a clearer view of the rest of the trajectory. Time is plotted as calendar time. A line at *R*_*e*_ = 1 is provided for convenience, as below this threshold the epidemic cannot be sustained. In grey is a heatmap of the inferred divergence times.

The results of our HSMRF-based model analysis are largely consistent with those of Vasylyeva *et al.* (2016), who also observed an increased rate of infection from 1995 to 2000 [62]. Our GMRF-based analysis, with its large decrease in *R*_*e*_(*t*), does not align with either the prevalence data or any analysis performed by Vasylyeva *et al.* (2016) [62]. While both our HSMRF-based and GMRF-based models estimated *R*_*e*_(*t*) *<* 1 throughout the 2000s, there is no evidence from HIV prevalence that the epidemic is decreasing [65, 66]. Examining the posterior distribution on phylogenies provides some insight into this apparent conflict: there are few infections inferred to have happened post-2000, and thus there is no information suggesting that *R*_*e*_(*t*) *>* 1 in this period. Previous coalescent analyses have favored a higher *R*_*e*_(*t*) persisting with no sign of a decrease, however such models can have difficulty inferring decreases in the absence of coalescent events [26]. This highlights the fact that while birth-death process and coalescent models are good at peering into the past, without birth (coalescent) events there is little to no information from which to infer birth (coalescent) rates and thus the posterior distribution is largely determined by the prior distribution. On the other hand, there is outside evidence that the epidemic has slowed since 2005 [67], so it is possible that our models are picking up on a real signal and simply exaggerating it.

### Code and data availability

We have implemented all models and necessary samplers in `RevBayes` [31]. Analysis were performed with `RevBayes` v1.0.11 (https://github.com/revbayes/revbayes). Scripts used for our real data analyses and simulations are available at github.com/afmagee/hsmrfbdp.

## Discussion and Conclusion

In this paper, we use a piecewise-constant birth-death model, combined with both GMRF and HSMRF prior distributions, to approximate arbitrary changes in both the birth and death rates through time. We implement these models in the statistical phylogenetic software platform `RevBayes`, allowing for both inference of birth-death process parameters using a phylogeny as data and for joint inference of BDP parameters, phylogeny, and nuisance parameters directly from molecular sequence data. Additionally, we present an intuitive scheme for setting the key hyperparameter for these models, the global shrinkage parameter, and provide an efficient and tuning-parameter free inference framework that enables inference for these high-dimensional models. We find that both GMRF- and HMRF-based models are capable of inferring variable birth rates and correctly rejecting variable models in favor of effectively constant models. When estimating birth rates, we see that in general the HSMRF-based model has higher precision than its GMRF counterpart, with little to no loss of accuracy. Applied to a macroevolutionary dataset of the Australian gecko family Pygopodidae (where birth rates are interpretable as speciation rates), our models detect a speciation-rate decrease in the last 12 million years. Applied to an infectious disease phylodynamic dataset of sequences from HIV subtype A in Russia and Ukraine, our models detect a complex pattern of variation in the rate of infection.

Through simulations we find that different functional forms of birth-rate variation produce unique challenges in estimating these forms, even if they share the same magnitude of change. Slow changes are easy to miss, intermediate shifts are largely detectable, and fast shifts are generally hard for the GMRF-based model but easy for the HSMRF-based model to estimate. It is likely that slow changes are difficult to detect because both priors prefer piecewise-constant trajectories to continuous variation. As there are relatively fewer births in the older part of the tree, the prior can more easily overwhelm the likelihood, leading to an effectively constant model being fit. This also likely contributes to the increase in uncertainty through time in the estimated rate. Fast shifts cause issues for the GMRF-based model because they require the global scale parameter *γ* to be large, which results in noisy and imprecise inference of slowly changing parts of the birth trajectory. At the same time, the GMRF-based model has a tendency to over-smooth the rapid changes. More recent changes are generally easier to detect than older changes, although very recent changes are often missed. Larger magnitude shifts are easier to detect than smaller magnitude shifts for both models, regardless of the functional form. In general, factors that make detecting shifts easier also exacerbate the poor behavior of the GMRF-based model. The HSMRF-based model often favors a trajectory with one or a few steeper shifts, even when the truth is a more gradual change. However, even if the duration of the shift is not accurately estimated, the HSMRF can recover the presence of rate variation even when the GMRF fails. Overall, we find that the performance of the HSMRF is quite good, and it is only clearly outperformed by the GMRF on a few types of birth rate trajectories.

Our simulations with time-varying death rates show that birth rates can be estimated quite well even in the face of difficulties inferring death rates. Serial sampling greatly improves the precision of the estimated birth rates and seems to mitigate the tendency of uncertainty to increase farther into the past. The increasing variability in the estimated rate is likely a function of both the increase in prior variability (due to the directional nature of our prior) and the reduced number of birth events in the past. The presence of serial samples can increase the number of observed birth events early in the process and thus improve both accuracy and precision in the estimated rate in older intervals. One theme that becomes even more evident in these simulations is that when the true birth-rate trajectory includes large jumps, the GMRF-based model will tend to infer spurious variability in the trajectory in regions where the birth rate should be small. When the death rate varies, the GMRF-based model is no more accurate than the HSMRF-based model in inferring the birth rate, but it infers substantially more variability than the HSMRF-based model, suggesting that much of this variability is in fact spurious. On the whole, though, we find that birth rates are generally well-estimated and that when there is serial sampling, the HSMRF-based model can capture the presence of variability in the death rate. This may at first seem contradictory with the findings of Louca and Pennell (2020) [68], who have shown that a fully-variable *λ*(*t*) and *μ*(*t*) cannot fully be identified from an extant phylogeny. For any tree there is an infinitely large “congruence class” of diversification-rate histories that are equally likely. We think that our results show the potential that priors provide for mitigating this problem. Bayesian inference introduces regularization to this problem in the form of prior distributions, which in general should reduce the size of the congruence class. Our results suggest that, at least in some instances, our priors are strong enough to ensure that there is only one set of birth- and death-rate trajectories that are plausible in light of observed data and imposed priors.

There are several avenues by which random field birth-death processes might be extended. It would be useful to devise models that can accurately infer slower declines, situations where the models we have put forth here have difficulty. One option for this would be to build second order Markov random field models, which can more easily collapse to linear models. These models have shown promise in coalescent modeling [29], but they have a higher risk of over-smoothing than first order models. Extending the models to include time-varying sampling rates may prove useful. Covariates may be added to time-varying birth-death models; previous work on birth-death models for macroevolution allowed for climate-dependent rates [20], while previous coalescent-based models considered the size of the region in which infections were found [30]. Adding covariates to the HSMRF-based model may allow for better success in inferring time-varying death rates by providing additional information. Models that allow for the serial sampling rate to vary may have better success (with or without covariates), as there is more direct information about this rate. However, cases where a number of samples have the same recorded age but there is not a sampling event (such as when some epidemiological sampling dates are available only to the year), may prove difficult. In such a case, the apparent variation in the sampling rate will likely overwhelm any signal of true variation in the sampling rate and may lead to erroneous estimates of the birth rate. Finally, for phylodynamic applications like HIV, it is clear that GMRF-based and HMRF-based birth-death models would benefit strongly from the inclusion of epidemiological data, which has been incorporated into time-homogenous birth-death-sampling-treatment process by Gupta *et al.* (2019) [69].

In this work, we have developed and explored the performance of an HSMRF-based birth-death model for time-varying birth and death rates. This model is capable of detecting slow or rapid shifts in birth rates, and can infer the timing of rapid shifts quite accurately. Detecting variability in birth and death rates simultaneously is problematic, but possible. The HSMRF-based models are is extensible, and incorporating covariates or variable sampling rates will widen the range of potential applications of these models. The GMRF-based models may also prove useful, but they have issues with both spurious inference of variability in birth rates and under-detection of variability in death rates. Therefore, we recommend using the HSMRF as Bayesian nonparametric priors for birth-death models.

## Acknowledgments

We are grateful to Arman Bilge, Jonathan Fintzi, Jean Feng, and Jared Grummer for helpful discussions, to James Faulkner for assistance with the MCMC samplers, and to Rebecca Magee for identifying a number of typos and inconsistencies in the manuscript. We also thank the reviewers and the guest editor for helpful suggestions that improved the manuscript. A.F.M. was supported by NSF grant DGE-1762114 and the ARCS Foundation Fellowship. S.H. was supported by the Deutsche Forschungsgemeinschaft (DFG) Emmy Noether-Program HO 6201/1-1. T.I.V. was supported by the Branco Weiss Fellowship and New College, University of Oxford. A.D.L. was supported by NSF grant DEB-1456098. V.N.M. was supported by the NIH grant R01-AI107034 and the NSF grant IIS-1561334.

