## Supplemental Materials/Appendices for "Locally adaptive Bayesian birth-death model successfully detects slow and rapid rate shifts"

### Supplementary Material

#### Simulating parameters

**Table S1.** The simulation values of the birth rates $\lambda_1$ and $\lambda_2$, and the times of change, $t_1$ and $t_2$, for all simulations. For the piecewise linear simulations, between $t_1$ and $t_2$, the birth rate is a linear interpolation between $\lambda_1$ and $\lambda_2$. For the piecewise constant simulations, there is only one time where the rate changes, $t_1$. For the constant-rate simulations, there are no change times and there is only one birth rate. The piecewise-linear simulations with $t_1 = t_2 = 50$ is also used when understanding the behavior of the models in the piecewise-constant scenarios, as it is nested in both sets of simulations (*). In all cases we round to 4 decimal places, the actual rates are not exactly equal for any pair of simulations.

| simulation type | fold change | $\lambda_1$ | $\lambda_2$ | $t_1$ | $t_2$ |
|---|---|---|---|---|---|
| piecewise linear* | 2 | 0.0727 | 0.0364 | 50 | 50 |
| piecewise linear | 2 | 0.0727 | 0.0364 | 37.5 | 62.5 |
| piecewise linear | 2 | 0.0727 | 0.0363 | 25 | 75 |
| piecewise linear | 2 | 0.0726 | 0.0363 | 12.5 | 87.5 |
| piecewise linear | 2 | 0.0726 | 0.0363 | 0 | 100 |
| piecewise constant | 2 | 0.0570 | 0.0285 | 10 | N/A |
| piecewise constant | 2 | 0.0603 | 0.0301 | 20 | N/A |
| piecewise constant | 2 | 0.0639 | 0.0320 | 30 | N/A |
| piecewise constant | 2 | 0.0681 | 0.0340 | 40 | N/A |
| piecewise constant | 2 | 0.0780 | 0.0390 | 60 | N/A |
| piecewise linear* | 4 | 0.0877 | 0.0219 | 50 | 50 |
| piecewise linear | 4 | 0.0877 | 0.0219 | 37.5 | 62.5 |
| piecewise linear | 4 | 0.0876 | 0.0219 | 25 | 75 |
| piecewise linear | 4 | 0.0876 | 0.0219 | 12.5 | 87.5 |
| piecewise linear | 4 | 0.0875 | 0.0219 | 0 | 100 |
| piecewise constant | 4 | 0.0586 | 0.0146 | 10 | N/A |
| piecewise constant | 4 | 0.0639 | 0.0160 | 20 | N/A |
| piecewise constant | 4 | 0.0703 | 0.0176 | 30 | N/A |
| piecewise constant | 4 | 0.0781 | 0.0195 | 40 | N/A |
| piecewise constant | 4 | 0.0999 | 0.0250 | 60 | N/A |
| constant | N/A | 0.0540 | N/A | XXX | N/A |

#### Performance on consant-rate datasets

To contextualize the performance of our models when the true scenario was constant-rate, we also inferred constant-rate birth-death processes on these trees. For constant-rate fits, the FC and RMASV performance measures are inapplicable, but the MAD and RP are still useful. Thus we compared the results of fitting constant-rate models to fitting GMRF-based and HSMRF-based models to constant-rate simulations, and present these in Figure S1. The GMRF-based model actually appears to perform better than the constant-rate model in terms of MAD, with an approximately 3% decrease in average MAD and an approximately 2% decrease in the 95th percentile of MAD. The HSMRF-based model performs trivially worse than a constant-rate model, with an approximately 1% worse average MAD and a 1% higher 95th percentile. Both models do suffer in precision, with the GMRF-based model's average RP approximately 31% higher than that of the constant-rate model and the HSMRF-based model's average RP 55% worse. As mentioned before, the cost of using a GMRF-based model or an HSMRF-based model instead of a constant-rate model is minor.

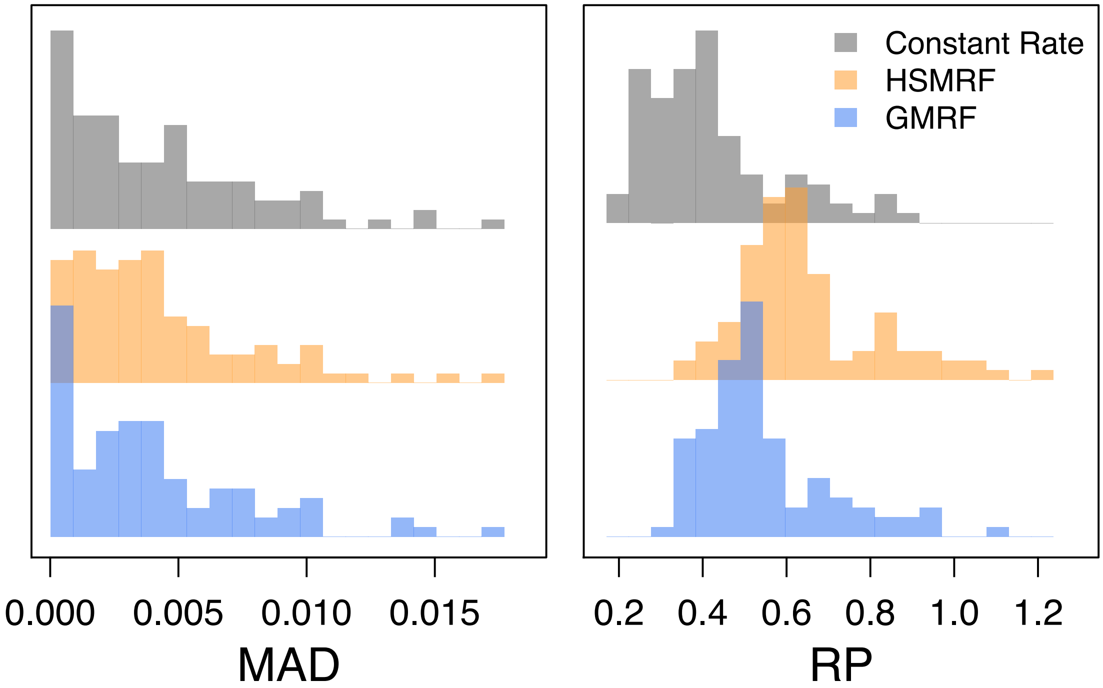


**Figure S1.** Performance of the models on simulated constant-rate datasets. MAD measures the error in the estimated trajectory. RP is a measure of precision, the average width of the 90% Credible Interval relative to the birth rate. We compare these measures between the using time-varying models (GMRF in blue, HSMRF in orange) for inference and using the true model (constant-rate, grey).

#### Additional examples of performance with time-varying birth rates

In the main text we showed examples of performance on a single constant-rate simulation and four time-varying trajectories. Here we present examples of performance for all of the time-varying birth rate simulations.

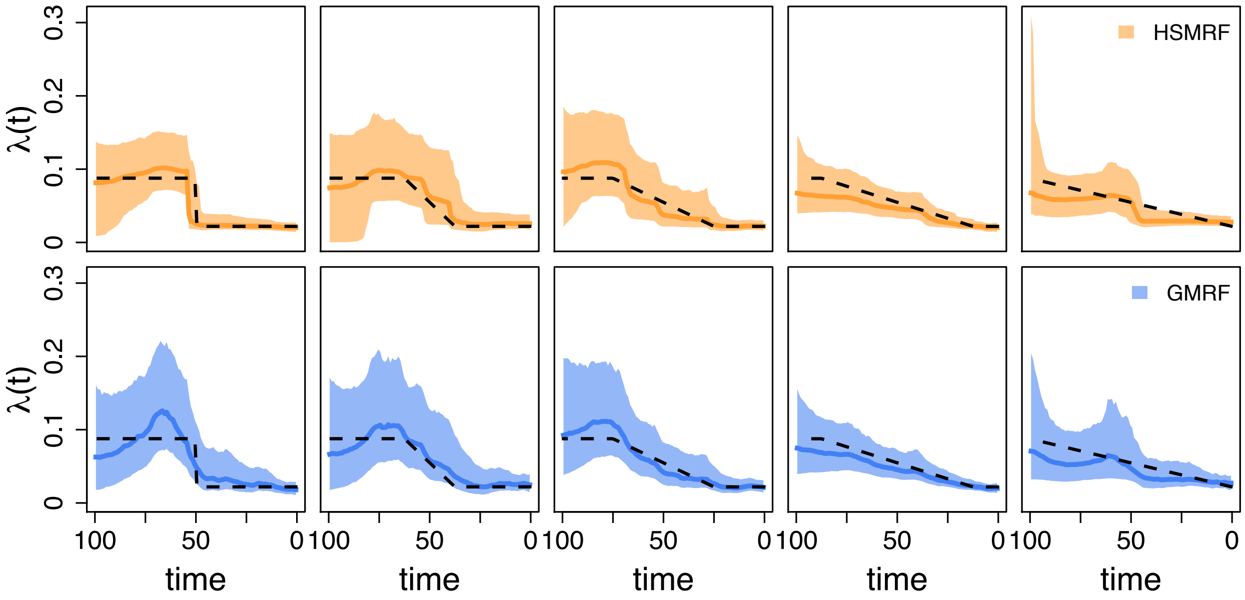


**Figure S2.** Inferred birth-rate trajectories from five individual simulations. The dashed line is the true simulating birth rate, the dark colored line is the posterior median trajectory (the median is taken separately for each grid cell), and the shaded region show the 90% Credible Interval (CI) for the rate. The columns demonstrate the effect of changing the shift duration (the length of the tree over which the birth rate changes), from an instantaneous shift to a constant change model. In each column, we show the simulation with the most average performance measured in terms of the Mean Absolute Deviation of both the GMRF and HSMRF.

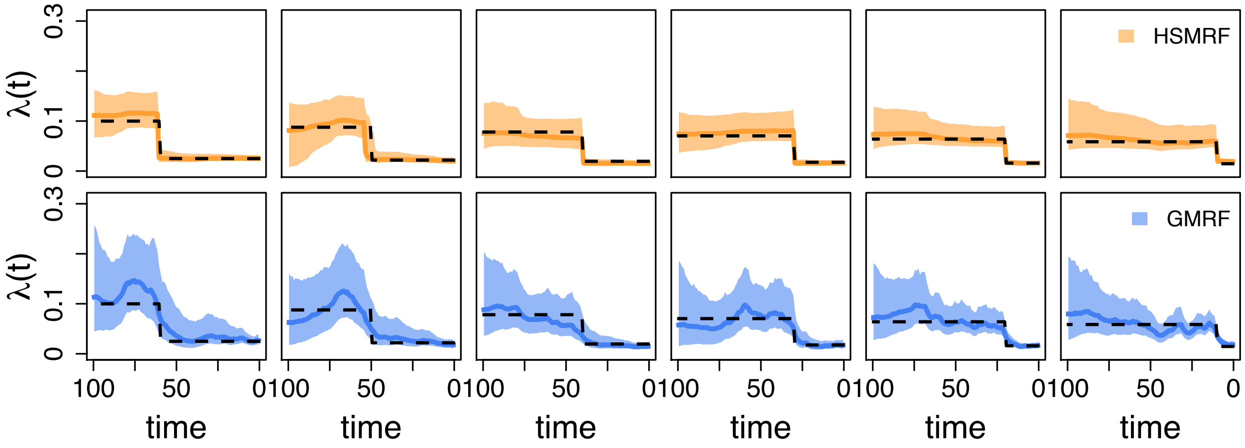


**Figure S3.** Inferred birth-rate trajectories from four individual simulations. The dashed line is the true simulating birth rate, the dark colored line is the posterior median trajectory (the median is taken separately for each grid cell), and the shaded region show the 90% Credible Interval (CI) for the rate. The columns demonstrate the effect of changing the location of the (instantaneous) shift from 60 time units before the present to 10 time units before the present. In each column, we show the simulation with the most average performance measured in terms of the Mean Absolute Deviation of both the GMRF and HSMRF.

#### Additional simulation results

In the main text we have focused on the results of simulations that had a four-fold net change. The results for two-fold changes are qualitatively similar, but here for completion we present versions of Figures 3 and 4 for two-fold shifts. We also present a complete table with all simulation parameters.

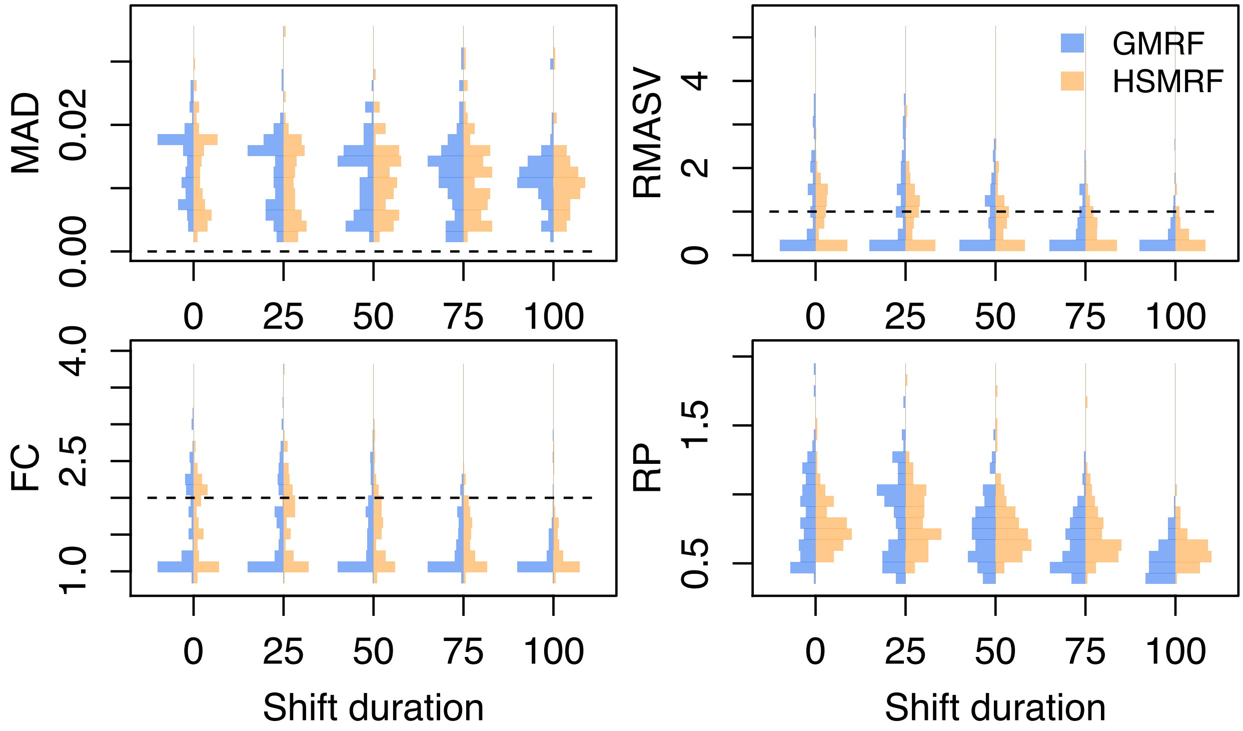


**Figure S4.** The effect of changing the duration of the two-fold rate shift, from instantaneous to the entire length of the trajectory. MAD measures the error in the estimated trajectory. RMASV measures the total amount of change relative to the true MASV, horizontal line at 1 for reference. FC measures the fold change from present to past, dotted line at true value for reference. RP is a measure of precision, the average width of the 90% Credible Interval relative to the birth rate.

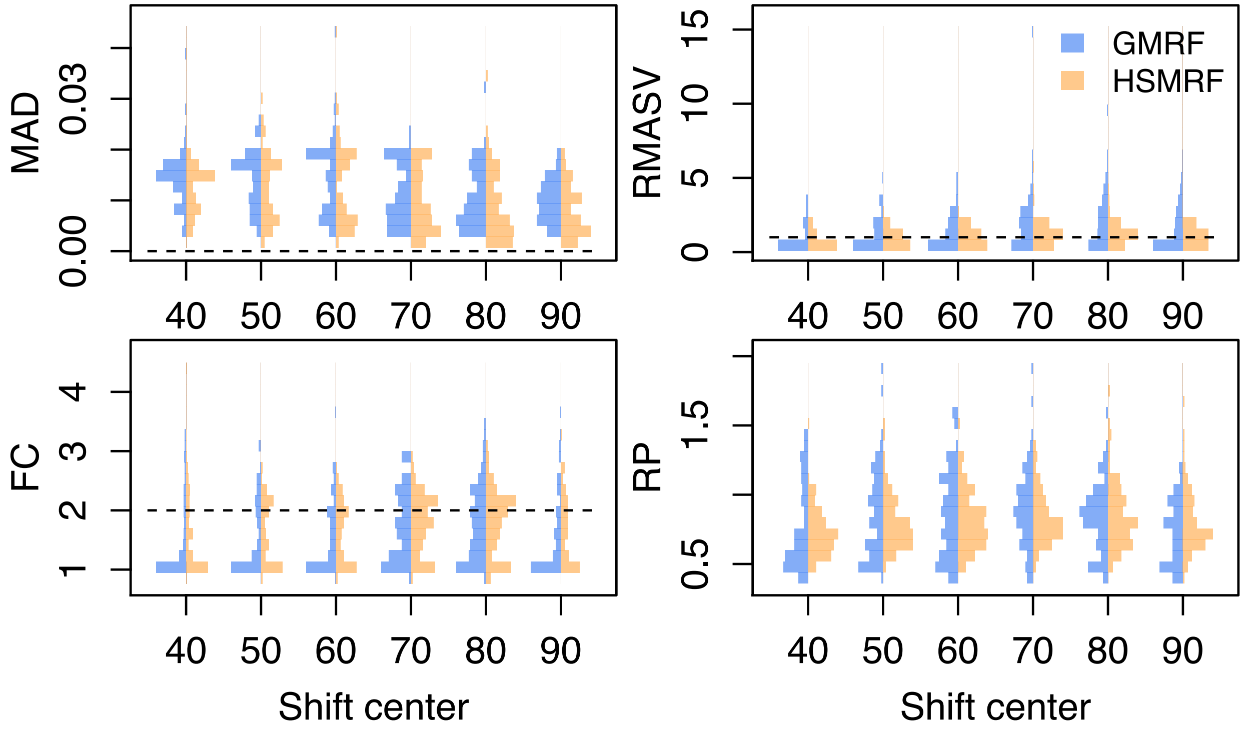


**Figure S5.** The effect of changing the location of the two-fold rate shift, from 60 to 10. MAD measures the error in the estimated trajectory. RMASV measures the total amount of change relative to the true MASV, horizontal line at 1 for reference. FC measures the fold change from present to past, dotted line at true value for reference. RP is a measure of precision, the average width of the 90% Credible Interval relative to the birth rate.

We lastly present a Figure for each experiment performed with additional summary measures. The additional summary measures are the minimum Effective Sample Size (minESS, taken across all values in the logfile, including parameters and transformed parameters), the coverage (coverage, percent of posterior birth rate 90% CIs containing the true value), and the Mean Squared Error (MSE).

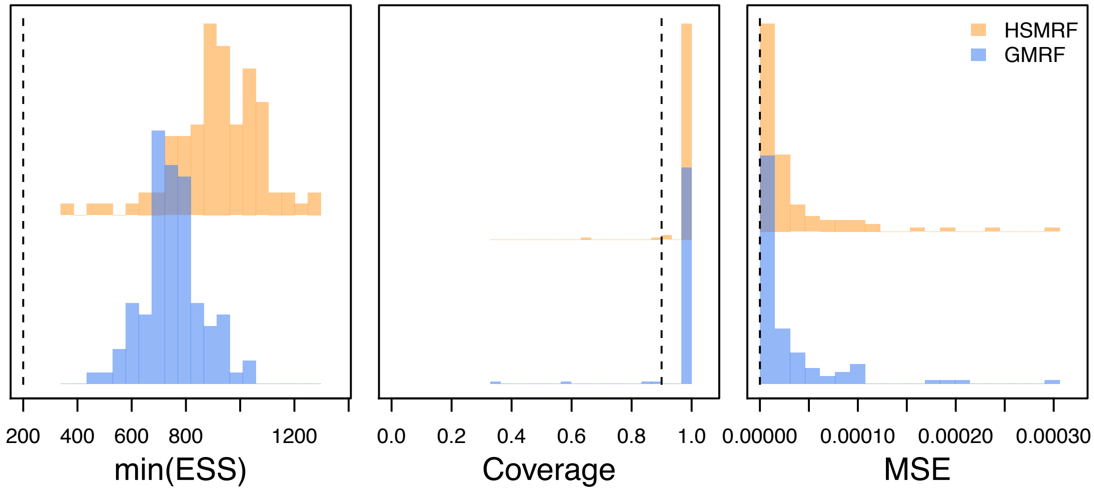


**Figure S6.** Performance on the constant-rate simulated datasets. minESS is the mininum ESS of all logged quantities (parameters, prior/likelihood/posterior, and transformed parameters), dashed line at 200 for reference. Coverage is the proportion of 90% CIs of the birth rates that include the true birth rate, dashed line at 0.9 for reference. MSE is the Mean Squared Error of the estimated trajectory, dashed line at 0 for reference.

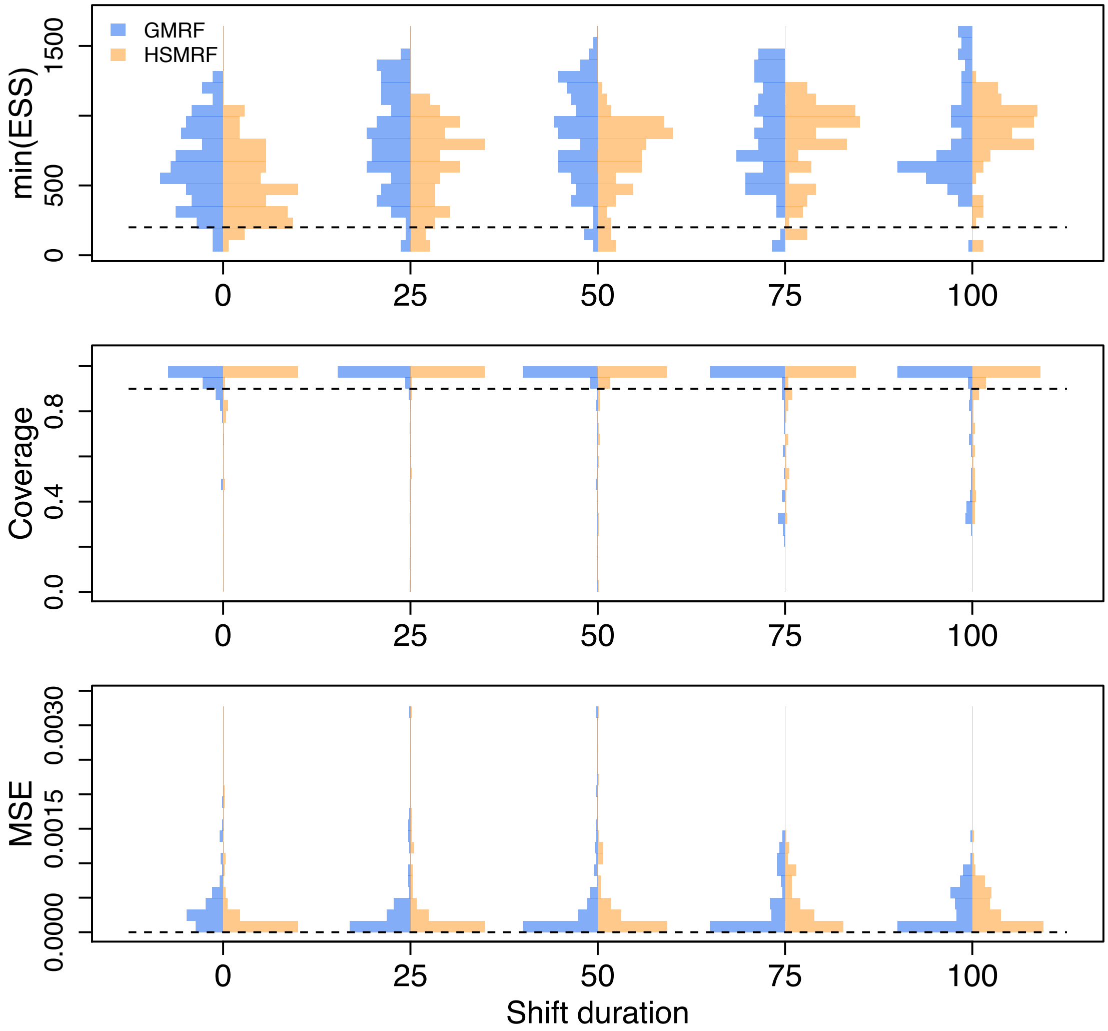


**Figure S7.** The effect of changing the duration of the four-fold rate shift, from instantaneous to the entire length of the trajectory. minESS is the mininum ESS of all logged quantities (parameters, prior/likelihood/posterior, and transformed parameters). Coverage is the percent of 90% CIs of the birth rates that include the true birth rate. MSE is the Mean Squared Error of the estimated trajectory.

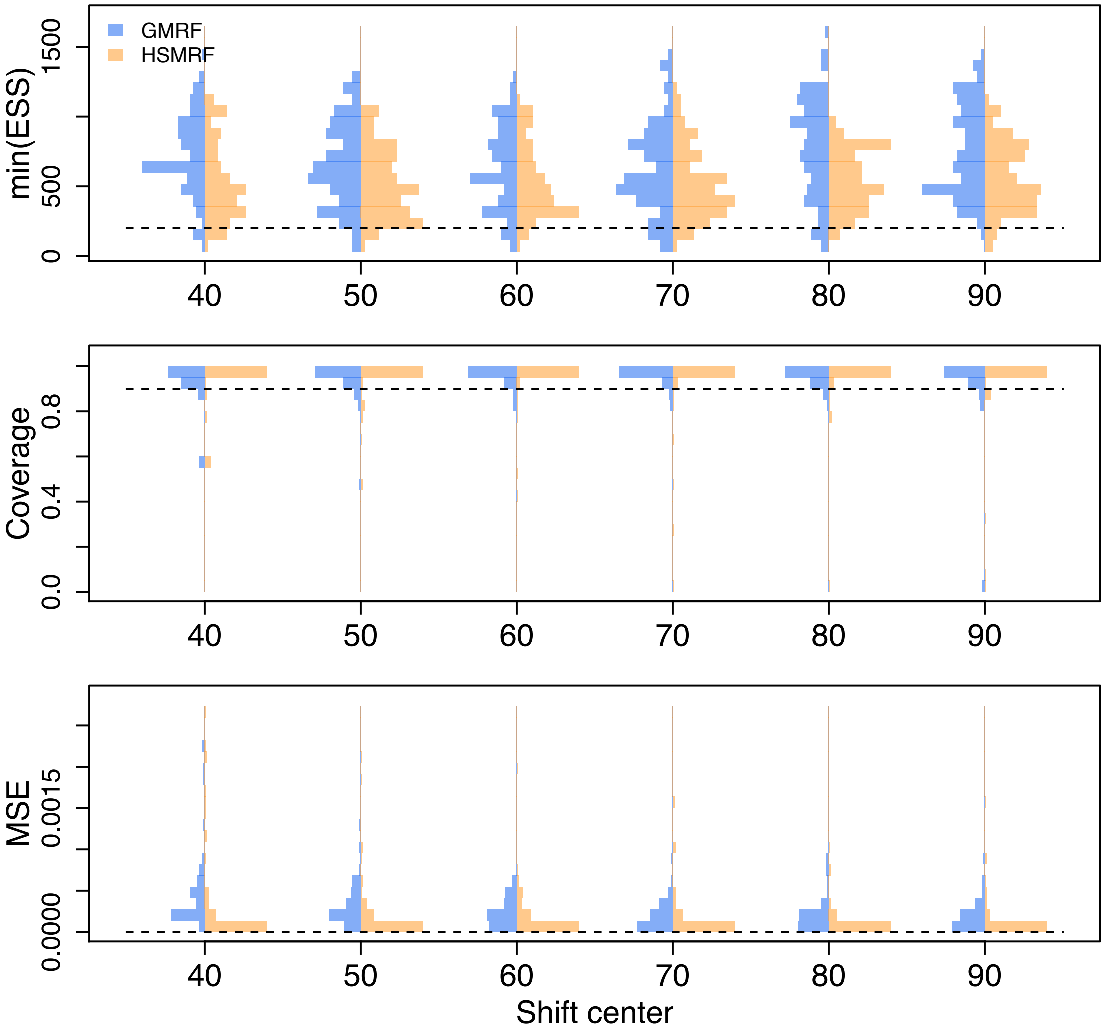


**Figure S8.** The effect of changing the location of the four-fold rate shift, from 60 to 10. minESS is the mininum ESS of all logged quantities (parameters, prior/likelihood/posterior, and transformed parameters). Coverage is the percent of 90% CIs of the birth rates that include the true birth rate. MSE is the Mean Squared Error of the estimated trajectory.

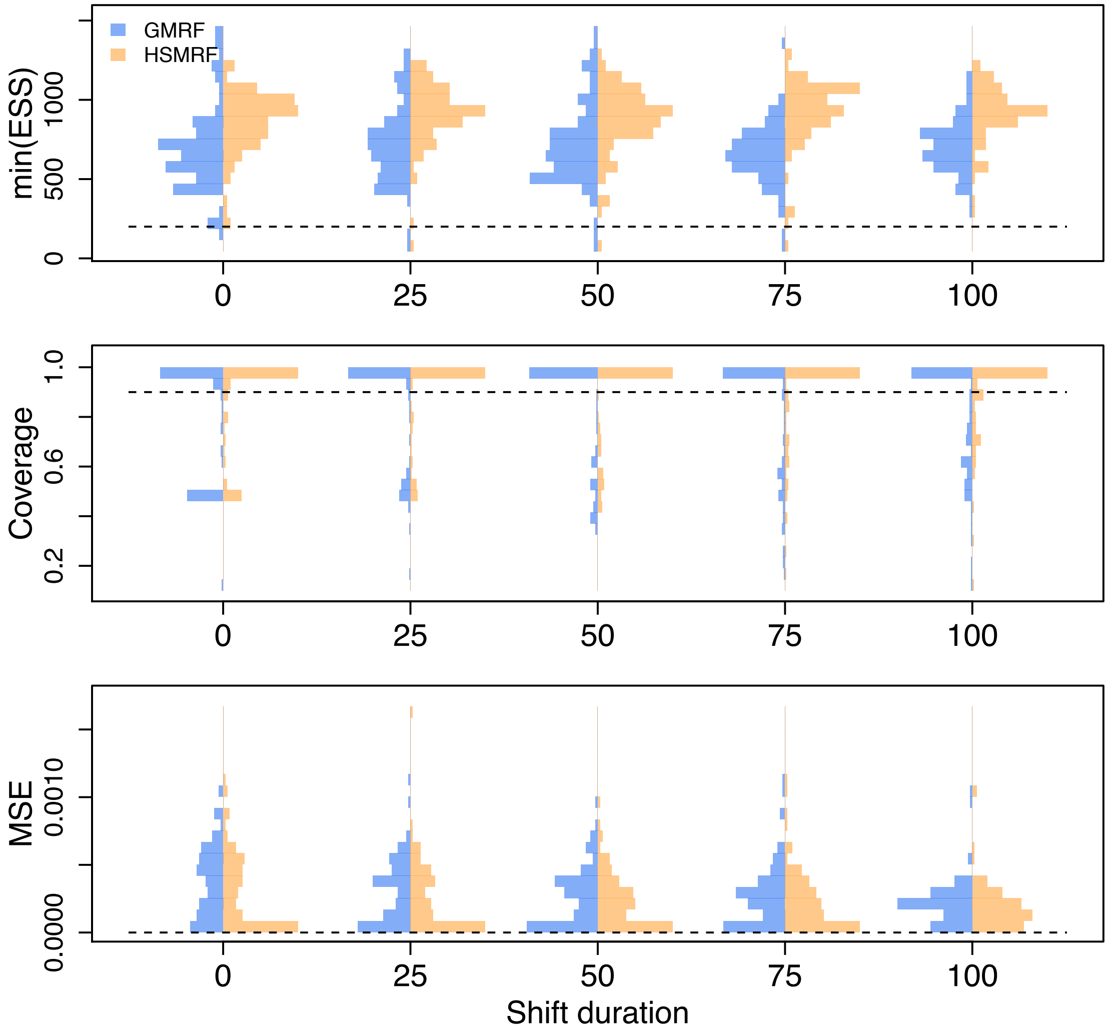


**Figure S9.** The effect of changing the duration of the two-fold rate shift, from instantaneous to the entire length of the trajectory. minESS is the mininum ESS of all logged quantities (parameters, prior/likelihood/posterior, and transformed parameters). Coverage is the percent of 90% CIs of the birth rates that include the true birth rate. MSE is the Mean Squared Error of the estimated trajectory.

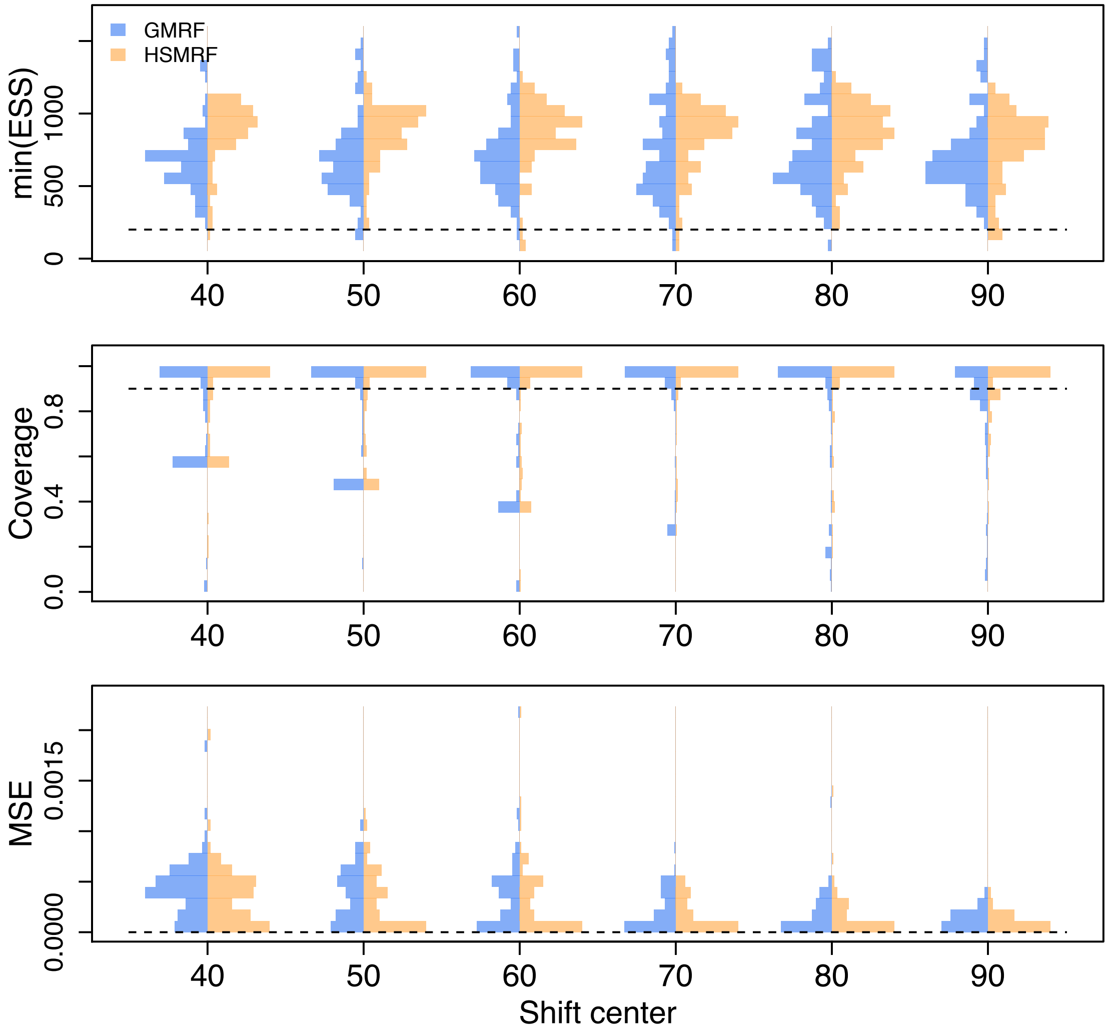


**Figure S10.** The effect of changing the location of the two-fold rate shift, from 60 to 10. minESS is the mininum ESS of all logged quantities (parameters, prior/likelihood/posterior, and transformed parameters). Coverage is the percent of 90% CIs of the birth rates that include the true birth rate. MSE is the Mean Squared Error of the estimated trajectory.

#### Estimating constant death rates

In the main text we showed examples of performance of estimating the death rate on two time-varying death-rate trajectories. Here we present examples of performance of estimating the death rate for all simulated scenarios where the true death rate was constant.

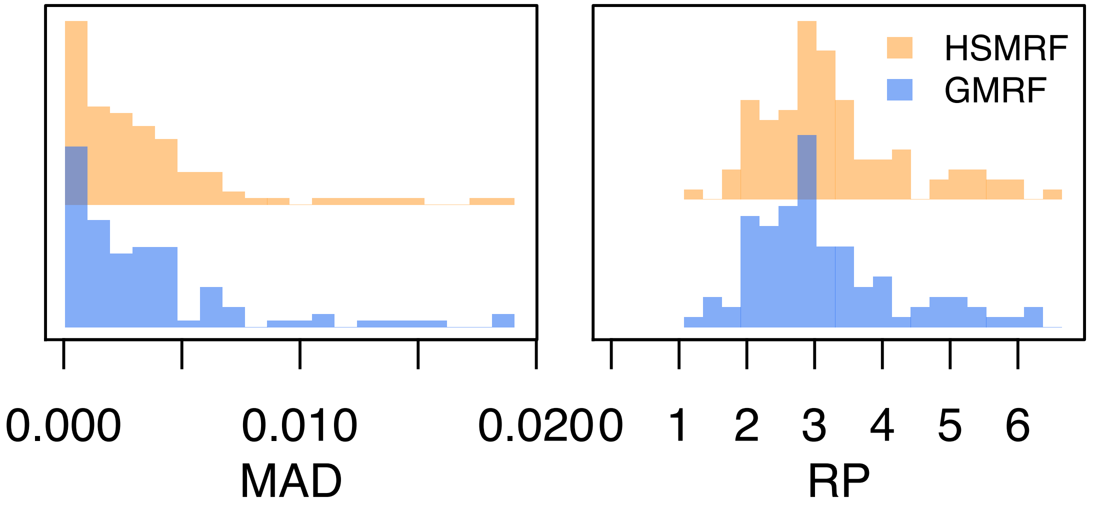


**Figure S11.** Performance of estimating the death rate in the constant-rate simulations. MAD measures the error in the estimated death rate. RP is a measure of precision, the width of the 90% Credible Interval relative to the death rate.

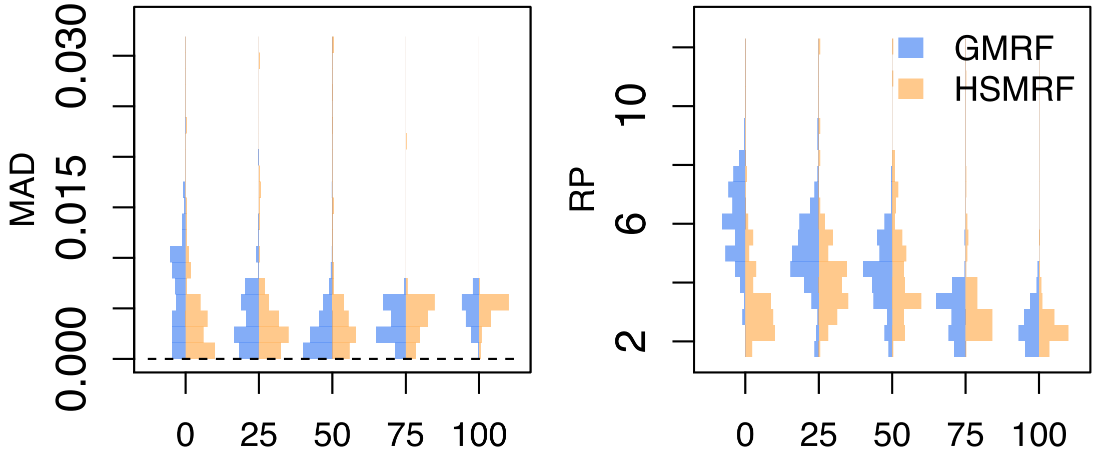


**Figure S12.** The effect of changing the duration of the four-fold rate shift, from instantaneous to the entire length of the trajectory on the estimated death rate. MAD measures the error in the estimated death rate. RP is a measure of precision, the width of the 90% Credible Interval relative to the death rate.

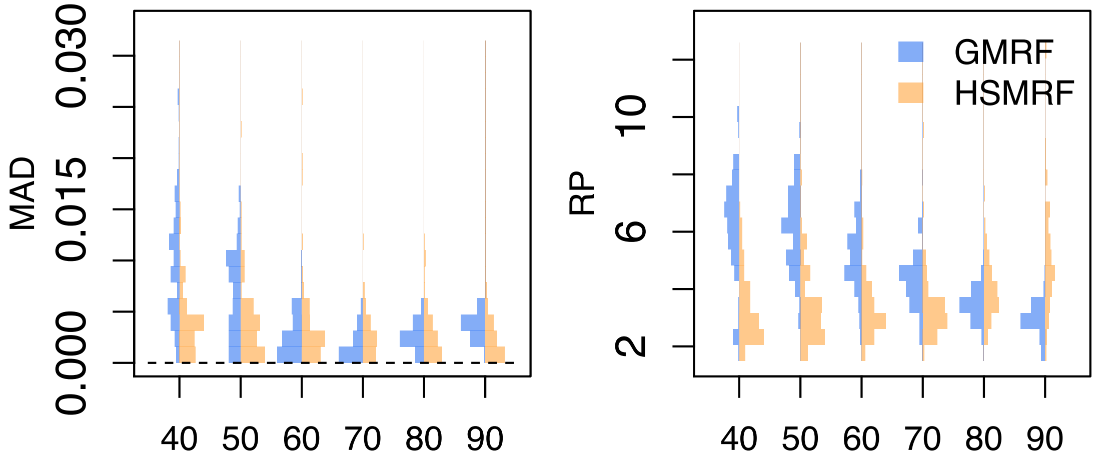


**Figure S13.** The effect of changing the location of the four-fold rate shift, from instantaneous to the entire length of the trajectory on the estimated death rate. MAD measures the error in the estimated death rate. RP is a measure of precision, the width of the 90% Credible Interval relative to the death rate.

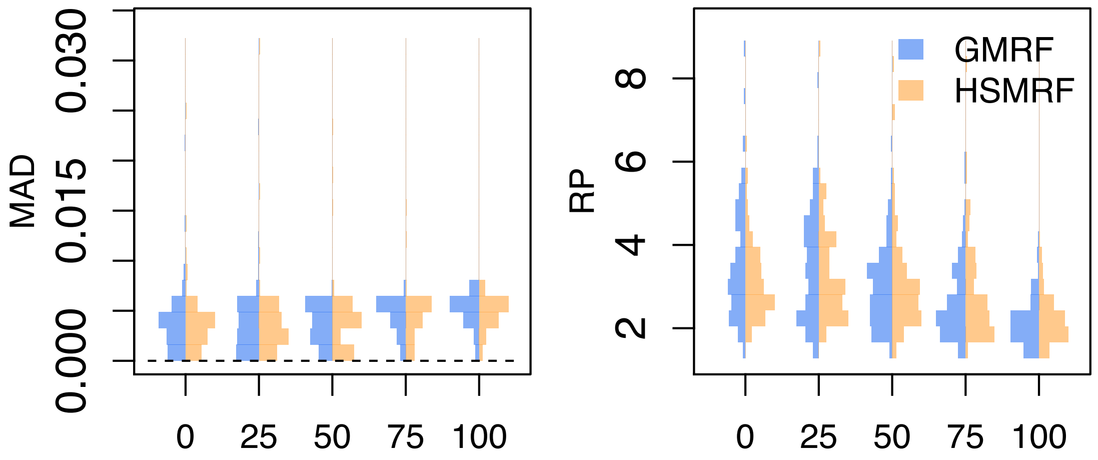


**Figure S14.** The effect of changing the duration of the two-fold rate shift, from instantaneous to the entire length of the trajectory on the estimated death rate. MAD measures the error in the estimated death rate. RP is a measure of precision, the width of the 90% Credible Interval relative to the death rate.

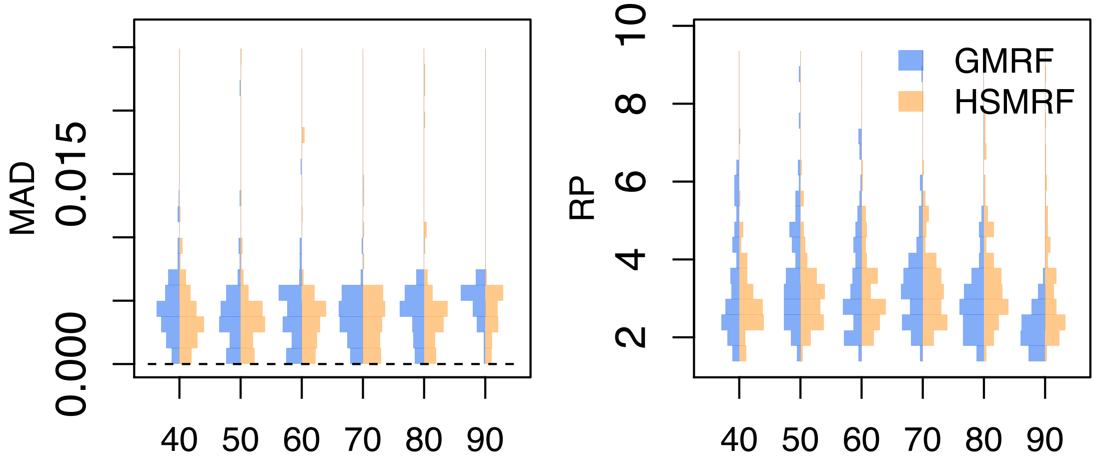


**Figure S15.** The effect of changing the location of the two-fold rate shift, from instantaneous to the entire length of the trajectory on the estimated death rate. MAD measures the error in the estimated death rate. RP is a measure of precision, the width of the 90% Credible Interval relative to the death rate.

#### Estimating time-varying death rates

In the main text we showed examples of performance when simulating from a model with time-varying death rates, and histograms of the MAD, RP, and RMASV performance measures. Here we present histograms with additional performance measures. We omit coverage, as the true death rate in many intervals is 0, making the coverage of the 90% CI seem artificially low. There is only one set of histograms for the minimum ESS as it is taken across all parameters in the posterior.

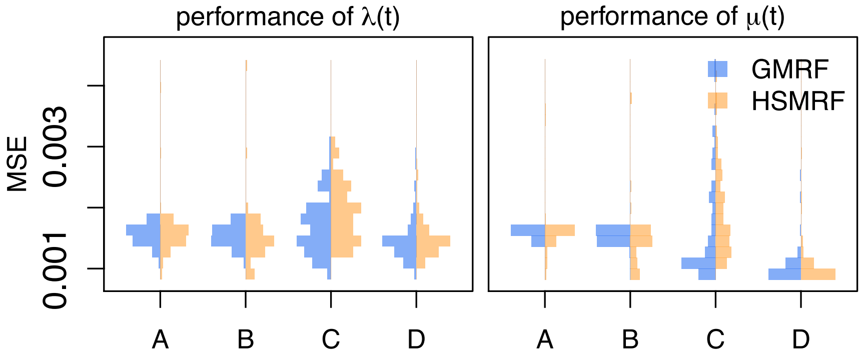


**Figure S16.** Performance of the models on simulated datasets with time-varying extinction. MSE is the Mean Squared Error of the estimated trajectory, dashed line at 0 for reference. The column labels A, B, C, and D identify the different combinations of tree simulations and analysis setup. A and B are analyses of trees with isochronous sampling, C and D heterochronous sampling. A and C are analyses where time-varying death rate, $\mu(t)$ is inferred, B and D where a constant death rate, $\mu$, is inferred.

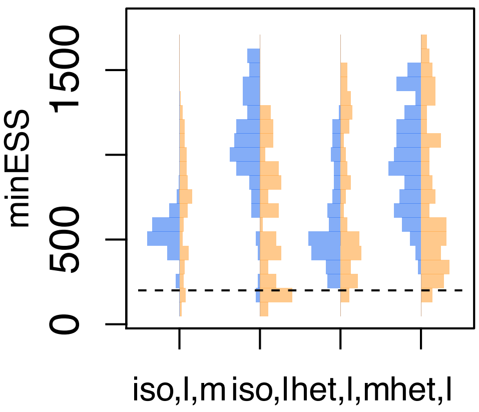


**Figure S17.** Performance of the models on simulated datasets with time-varying extinction. minESS is the mininum ESS of all logged quantities (parameters, prior/likelihood/posterior, and transformed parameters), dashed line at 200 for reference. The column labels A, B, C, and D identify the different combinations of tree simulations and analysis setup. A and B are analyses of trees with isochronous sampling, C and D heterochronous sampling. A and C are analyses where time-varying death rate, $\mu(t)$ is inferred, B and D where a constant death rate, $\mu$, is inferred.

#### Non-centered parameterizations

Markov random fields can be parameterized by the distribution of the change between neighbors, $\Delta_i := \lambda^*_{i+1} - \lambda^*_i$. In a GMRF, these "increments" of the field, are iid Normal(0,$\gamma$) random variables, and we can obtain $\lambda_i$ from $\lambda_1, \lambda^*_2, \dots, \lambda^*_{i-1}$ as $\lambda_i = \lambda_1 \exp(\sum_1^{n-1} \Delta_i)$. This approach is called the non-centered parameterization, and simplifies MCMC sampling. The usefulness of non-centralized parameterizations is that they break the dependency between adjacent field parameters by making every move to field parameters multivariate. A proposed change to $\lambda^*_i$ must fit within the prior imposed by $\lambda^*_{i-1}$ and must place a reasonable prior on $\lambda^*_{i+1}$. However, a change to $\Delta_i$ implicitly changes not only $\lambda^*_i$, but $\lambda^*_i, \dots, \lambda^*_n$ as well.

We can circumvent directly placing a half-Cauchy(0,$\zeta$) prior on $\gamma$, by placing a half-Cauchy(0,1) prior on $\gamma$ and using $\gamma\zeta$ in its place. This rescaling enables us to use a Gibbs sampler for $\gamma$ (for more details on this sampler see the next section), which enables us to appropriately explore the tails of the halfCauchy distribution where standard Metropolis Hastings moves fail. Thus, in our non-centered, rescaled parameterization of the GMRF, the increments are given by $\Delta_i \sim \text{Normal}(0, \gamma^2\zeta^2)$. For HSMRFs we expand this re-scaling to the $\sigma_i$, and rewrite $\sigma_i \sim \text{halfCauchy}(0, \gamma)$ as $\sigma_i \sim \text{halfCauchy}(0, 1)$ and where we would use $\sigma_i$ we use $\sigma_i\gamma\zeta$. For our non-centered, rescaled HSMRF, the increments are given by $\Delta_i \sim \text{Normal}(0, \sigma_i^2\gamma^2\zeta^2)$. We show the models as DAGs in Figure S18 as they are setup in our `RevBayes` analyses for full joint analysis of phylogeny and birth rates.

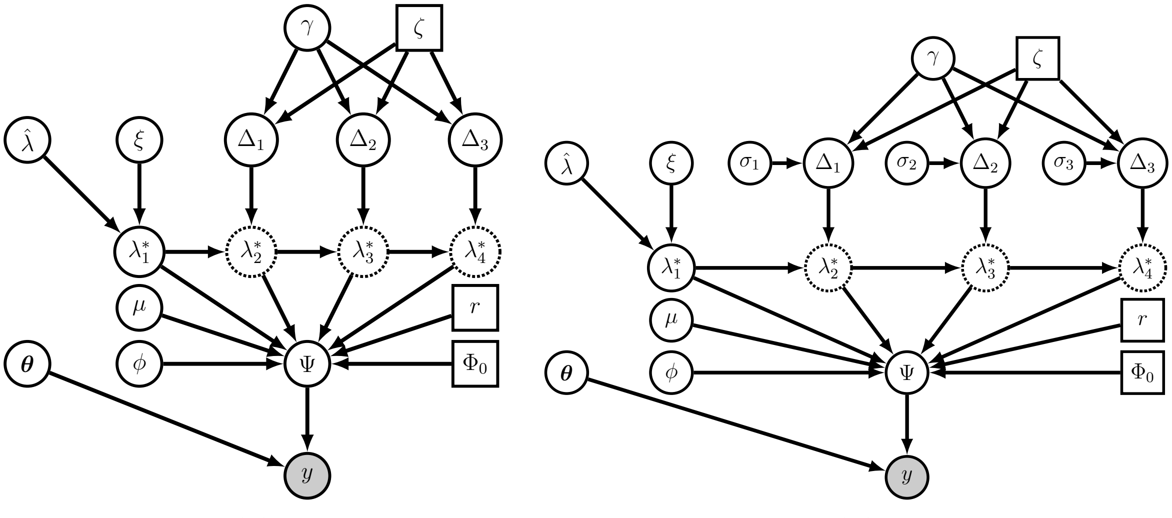


**Figure S18.** The parameterization of our models as setup in `RevBayes`, shown as a grid of size 4 for simplicity. In (a) we show our GMRF-based model, and in (b) our HSMRF-based model. The use of $\boldsymbol{\Delta}$ as parameters instead of $\boldsymbol{\lambda}^*_{2:n}$ simplifies MCMC by allowing us to sample independent variables. The rescaling of the $\boldsymbol{\Delta}$ by separating out $\boldsymbol{\sigma}$, $\gamma$, and $\zeta$ is required for the Gibbs sampler to run, but also serves to minimize the number of layers in the model. In addition to $\boldsymbol{\lambda}$, the tree prior is specified by the serial sampling rate $\phi$, death rate $\mu$, sampling probability at the present $\Phi_0$, and conditional probability of death upon sampling (treatment probability) $r$. We place all substitution and clock model parameters in $\boldsymbol{\theta}$, such that given $\boldsymbol{\theta}$ and the tree $\Psi$, the likelihood of the data $\boldsymbol{D}$ can be computed. When drawing the model as a DAG, squares represent constant values, closed circles stochastic values, open circles deterministic transformations of other nodes, and shaded circles observed stochastic values (data).

#### MCMC procedures

The major drawback to shrinkage priors is that the same fat tailed behavior that makes them adaptive makes inference difficult. We find that sampling is best accomplished under the non-centered, rescaled parameterization. However, while these model reparameterizations make sampling easier, basic Metropolis-Hastings MCMC proposals proved incapable of exploring the tails of the posterior. Our solution is to use an elliptical slice update to all increments of the random fields, $\boldsymbol{\Delta}$, Gibbs updates to all field hyperparameters, $\boldsymbol{\sigma}$ and $\gamma$, and standard MH updates to $\lambda^*_1$, $\mu$, and any substitution or tree model parameters.

Elliptical slice sampling [44] exploits the geometry of multivariate normal priors. It slices not in a straight line but along an ellipse that should maintain higher posterior probability. Mechanistically, this is accomplished by drawing proposed values from the multivariate normal prior and searching for an angle of rotation $\theta$ (to combine the proposed and current values) that is acceptable under the likelihood. In our case, the $\boldsymbol{\Delta}$ are all independently distributed, thus we can circumvent the usual need for matrix decomposition when drawing the proposed values, decreasing the time it takes to run the sampler. In Algorithm 1 we provide pseudocode for this special case, for a full explanation of the general case we refer readers to Murray *et al.* [44]. We note that in our implementation, there are two differences from the algorithm as presented in Algorithm 1, tuning and a hard limit on iterations. We allow for tuning by initializing $\theta \sim \text{Uniform}(0, m)$ ($m \leq 2\pi$) and tuning $m$ so as to reduce

**Algorithm 1** Elliptical Slice Sampler

**Input:** $\boldsymbol{\Delta}^t$ the value of $\boldsymbol{\Delta}$ at the current MCMC step, $\boldsymbol{\sigma}^t$ the standard deviations of $\boldsymbol{\Delta}^t$, function $f(x)$ that returns the (unnormalized) log-probability density for the downstream DAG for $\boldsymbol{\Delta} = \boldsymbol{x}$, $n$ the size of $\boldsymbol{\Delta}$.
Initialization:
draw $u \sim \text{Uniform}(0, 1)$
set $y \leftarrow f(\boldsymbol{\Delta}^t) + \ln(u)$
**for** i in 1:n **do**
  draw $\nu_i \sim \text{Normal}(0, (\sigma_i^t)^2)$
**end for**
draw $\theta \sim \text{Uniform}(0, 2\pi)$
set $L \leftarrow \theta - 2\pi$
set $R \leftarrow \theta$
**for** i in 1:n **do**
  set $x_i \leftarrow \Delta_i^t \cos(\theta) + \nu_i \sin(\theta)$
**end for**
Run sampler:
**while** $f(x) \leq y$ **do**
  **if** $\theta > 0$ **then**
    $R \leftarrow \theta$
  **else**
    $L \leftarrow \theta$
  **end if**
  draw $\theta \sim \text{Uniform}(L, R)$
  **for** i in 1:n **do**
    set $x_i \leftarrow \Delta_i^t \cos(\theta) + \nu_i \sin(\theta)$
  **end for**
**end while**
set $\boldsymbol{\Delta}^{t+1} = \boldsymbol{x}$

the total number of iterations required to accept. Larger $m$ correspond to bolder moves, and when the data are particularly informative, most moves will not be accepted until $\theta$ is close to 0, thus starting with a smaller $m$ reduces the number of iterations of the loop. We also cap the number of loop iterations at 2000, at which point the sampler will abort or, if the user desires, accept a move of $\theta = 0$ (after 2000 iterations, $\theta \approx 0$).

Gibbs sampling has been proposed for horseshoe distributions in regression contexts for all parameters in the model [70]. Faulkner *et al.* (2018) have expanded this to work on arbitrary HSMRF models and GMRF models [29]. This Gibbs sampling approach nominally requires further reformulating the model, beyond the parameterization discussed above. Specifically, it reparameterizes halfCauchy variables as mixtures of inverseGamma variables by adding yet another layer of auxiliary variables. To make implementing our models more user-friendly (by reducing the number of hierarchical layers in the model), we implement this Gibbs sampler in `RevBayes` to operate directly on the non-centered parameterization we have described thus far. This is permissible because the conditional distributions of the new auxiliary variables only depend on the values of $\sigma_i$ and $\gamma$. Thus they can be drawn at every sampling step conditioned on the current values, and then new values of $\sigma_i$ and $\gamma$ can

**Algorithm 2** Gibbs sampler for HSMRF

**Input:** $\mathbf{\Delta}^t$ the value of $\mathbf{\Delta}$ at the current MCMC step, $\boldsymbol{\sigma}^t$ the standard deviations of $\mathbf{\Delta}^t$, $\gamma^t$ the global scale parameter, $\zeta$ the global scale hyperparameter, $n$ the size of $\mathbf{\Delta}$.
**for** i in 1:n **do**
  draw $\psi_i \sim \text{InverseGamma}(1, 1 + 1/\sigma_i^2)$
  draw $(\sigma_i^{t+1})^2 \sim \text{InverseGamma}(1, \psi_i^{-1} + (\Delta_i^t)^2/(2(\gamma^t)^2\zeta^2))$
**end for**
draw $\xi \sim \text{InverseGamma}(1, 1 + 1/(\gamma^t)^2)$
draw $(\gamma^{t+1})^2 \sim \text{InverseGamma}(1, \xi^{-1} + 1/(2\zeta^2)\sum_{i=1}^n \Delta_i^2/\sigma_i^2)$

**Algorithm 3** Gibbs sampler for GMRF

**Input:** $\mathbf{\Delta}^t$ the value of $\mathbf{\Delta}$ at the current MCMC step, $\gamma^t$ the global scale parameter, $\zeta$ the global scale hyperparameter, $n$ the size of $\mathbf{\Delta}$.
draw $\xi \sim \text{InverseGamma}(1, 1 + 1/(\gamma^t)^2)$
draw $(\gamma^{t+1})^2 \sim \text{InverseGamma}(1, \xi^{-1} + 1/(2\zeta^2)\sum_{i=1}^n \Delta_i^2)$

be drawn and returned, leaving the auxiliary variables entirely behind the scenes. For the HSMRF-based model, the Gibbs sampler requires specifying a value of $\zeta$, and the rest of the non-centered HSMRF model is as follows,

$$\gamma \sim \text{halfCauchy}(0, 1),$$
$$\sigma_i \sim \text{halfCauchy}(0, 1),$$
$$\Delta_i \mid (\sigma_i, \gamma, \zeta) \sim \text{Normal}(0, \sigma_i^2\gamma^2\zeta^2).$$

For the GMRF-based model, the Gibbs sampler requires specifying a value of $\zeta$, and the rest of the non-centered GMRF model is as follows,

$$\gamma \sim \text{halfCauchy}(0, 1),$$
$$\Delta_i \mid (\gamma, \zeta) \sim \text{Normal}(0, \gamma^2\zeta^2).$$

We present pseudocode for implementing the Gibbs samplers in Algorithms 2 and 3, noting that the samplers operate on $\gamma^2$ and $\boldsymbol{\sigma}^2$, whereas our parameterization is on $\gamma$ and $\boldsymbol{\sigma}$. In practice inverseGamma variates are obtained by drawing Gamma random variables and inverting them. Thus in our implementation of the algorithms we draw $\boldsymbol{\psi}^{-1}$, $\xi^{-1}$, $\boldsymbol{\sigma}^{-2}$, $\gamma^{-2}$ directly from gamma distributions and invert variables only as needed.

#### Diagnosing MCMC convergence

When performing Bayesian analysis of models, performance diagnostics are absolutely necessary. We use the Potential Scale Reduction Factor (PSRF) to determine if two chains have converged [71]. We follow Vehtari *et al.* (2019) in using rank-transformed variables in our PSRF calculations to avoid issues induced by fat-tailed posterior distributions because the original PSRF assumes normally distributed variables [72]. For each variable in our posterior distribution we compute two convergence diagnostics, the PSRF of the rank-transformed variables, $\hat{R}_r$, and the PSRF of the rank-transformed folded variables, $\hat{R}_{rf}$, which can capture differences in variance between chains. To ensure that all our summaries are based on trustworthy MCMC runs, we discard the entire posterior for any analysis where

$\max(\hat{R}_r, \hat{R}_{rf}) > 1 + \epsilon$. In the simulations with a constant death rate, a cutoff using $\epsilon = 0.01$ rarely results in more than 2 or 3 analyses discarded per hundred analyses (HSMRF or GMRF analyses of simulated datasets, all constant-rate analysese passed). In the simulations with a time-varying death rate, this cutoff generally results in over 15 analyses being discarded per 100 analyses. Using $\epsilon = 0.02$ results instead in a maximum of 20 analyses discarded, and qualitatively similar results figures. We thus use $\epsilon = 0.01$ for the constant death simulations and $\epsilon = 0.02$ for the variable death simulations. We also calculate the effective sample size (ESS) for all parameters for the two chains combined for each model, and find that almost all parameters have ESS $> 200$. ESS is generally larger for the time-varying death rate simulations, which were run twice as long (and thinned twice as much, 400,000 sampled every 400 instead of 200,000 sampled every 200).

#### The size of the grid

In all of our simulations and analyses, we have employed a grid with 100 cells (a grid of size 100). Though performance with a grid of size 100 appears to be good, some may wonder if this grid is sufficiently large, or even too large. Using a tree sampled from the posterior distribution for the Pygopodidae HSMRF analysis, we here address that question. We take this tree to be data, and run analyses on it with grid sizes of 10,20,50,100, and 200, which we plot in Figure S19. We run two independent chains per analaysis, and check rank-based PSRF and ESS for all parameters in both GMRF and HSMRF analyses of all 5 grid sizes. The minimum ESS was 176 and the maximum PSRF was 1.016, indicating adequate convergence for our purposes. While the inferences using smaller grids (10 or 20) look rather different from the others, it is visually hard to distinguish the inferences using larger grids (50, 100, and 200). Thus, 100 is a reasonable grid size, in that it produces inferences similar to larger grids, while being faster to run.

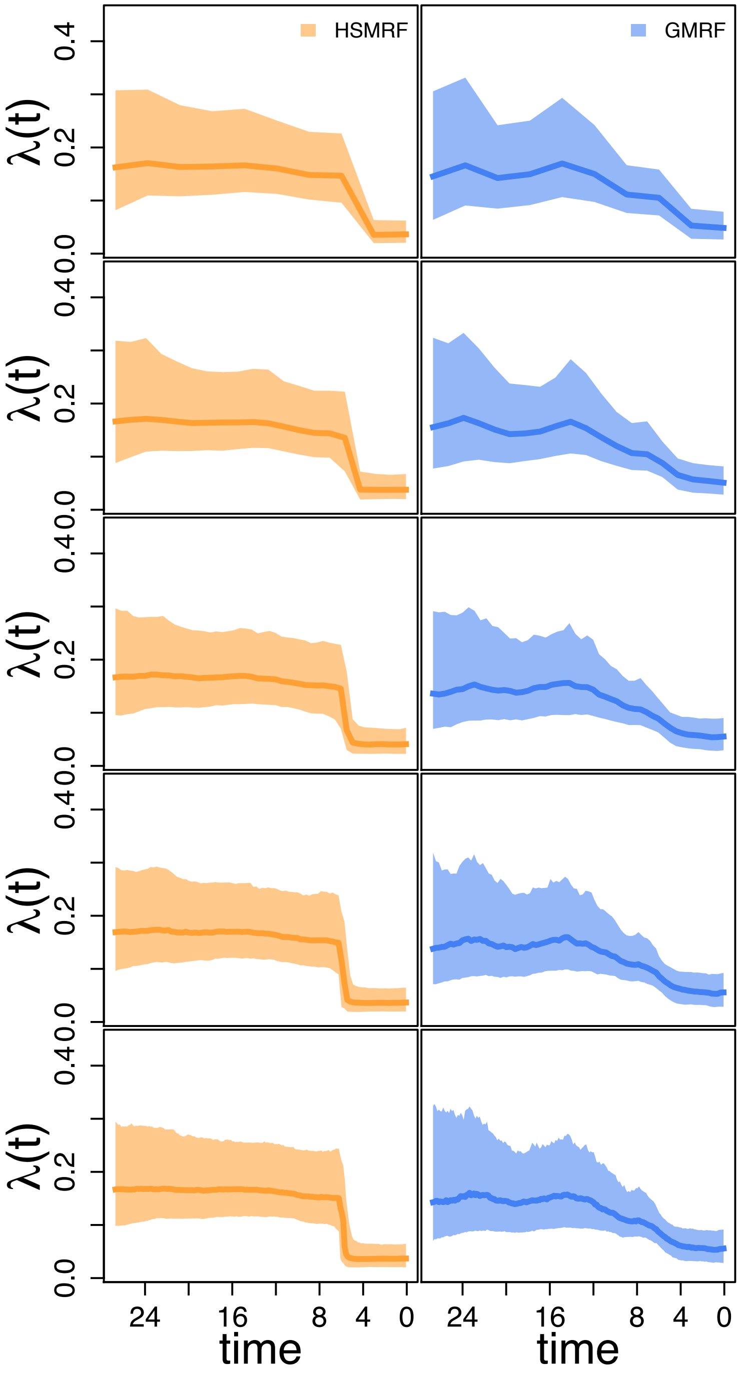


**Figure S19.** The effect of the size of the grid on estimated speciation rates in Pygopodidae. The tree is fixed to a tree from the posterior distribution from the Pygopodidae HSMRF analysis. From top to bottom, the grid sizes used for inference are 10, 20, 50, 100, and 200.

#### Sensitivity to death-rate prior

In all of our analyses of simulated data, and our empirical analysis of Pygopodidae, we employed an Exponential(10) prior on the death (extinction) rate. However, it is plausible that there is some prior sensitivity to the extinction rate prior. Using a tree sampled from the posterior distribution for the Pygopodidae HSMRF analysis (the same as with the grid size analyses), we here address that question. We take this tree to be data, and run analyses on it with several different death-rate priors. In all cases we hold the birth-rate prior constant and use the same prior we have used everywhere. Three of these are exponential priors, with rates of 1, 10, and 100. Two of these priors are empirical Bayes priors, where we fit a constant-rate birth-death model to the dataset first, and then using the method of moments fit either a Lognormal or a Gamma distribution to the posterior. Following May *et al.* [22], we inflate the variance 10-fold. These empirical Bayes priors have mean 0.011 and variance 0.0013. We run two independent chains per analaysis, and check rank-based PSRF and ESS for all parameters in both GMRF and HSMRF analyses of all 5 analyses. The minimum ESS was 270 and the maximum PSRF was 1.018, indicating adequate convergence for our purposes.

The estimated death rate is notably sensitive to the prior (Figure S20). The Exponential(1) and Exponential(10) priors produce the most similar posterior distributions, and appear to exert the least influence. The estimated birth rates are much less sensitive to the prior on the death-rate S21. The credible intervals are wider for the Exponential(1) and Exponential(10) priors than the other three priors. The magnitude of the shift is also somewhat different between the Exponential(1) and Exponential(10) priors (FC of approximately 4.7 for the HSMRF analyses and 2.6 for the GMRF analyses) and the other three priors (FC of approximately 3.9 for the HSMRF analyses and 1.9 for the GMRF analyses). However, qualitatively all trajectories are very similar and show clear evidence of a shift at approximately 6 Ma. Thus, even if the exact estimates of the birth rate change somewhat with the estimated death rate, the pattern remains consistent.

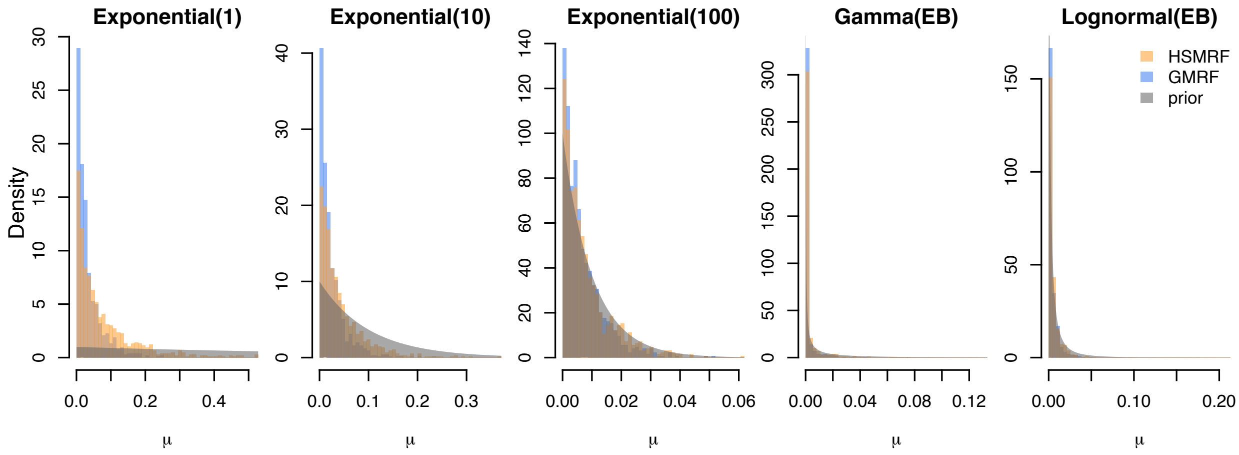


**Figure S20.** The effect of the death-rate prior on the estimated death rate. The tree is fixed to a tree from the posterior distribution from the Pygopodidae HSMRF analysis. The plots are not scaled consistently in order to facilitate prior-posterior comparisons for each prior individually.

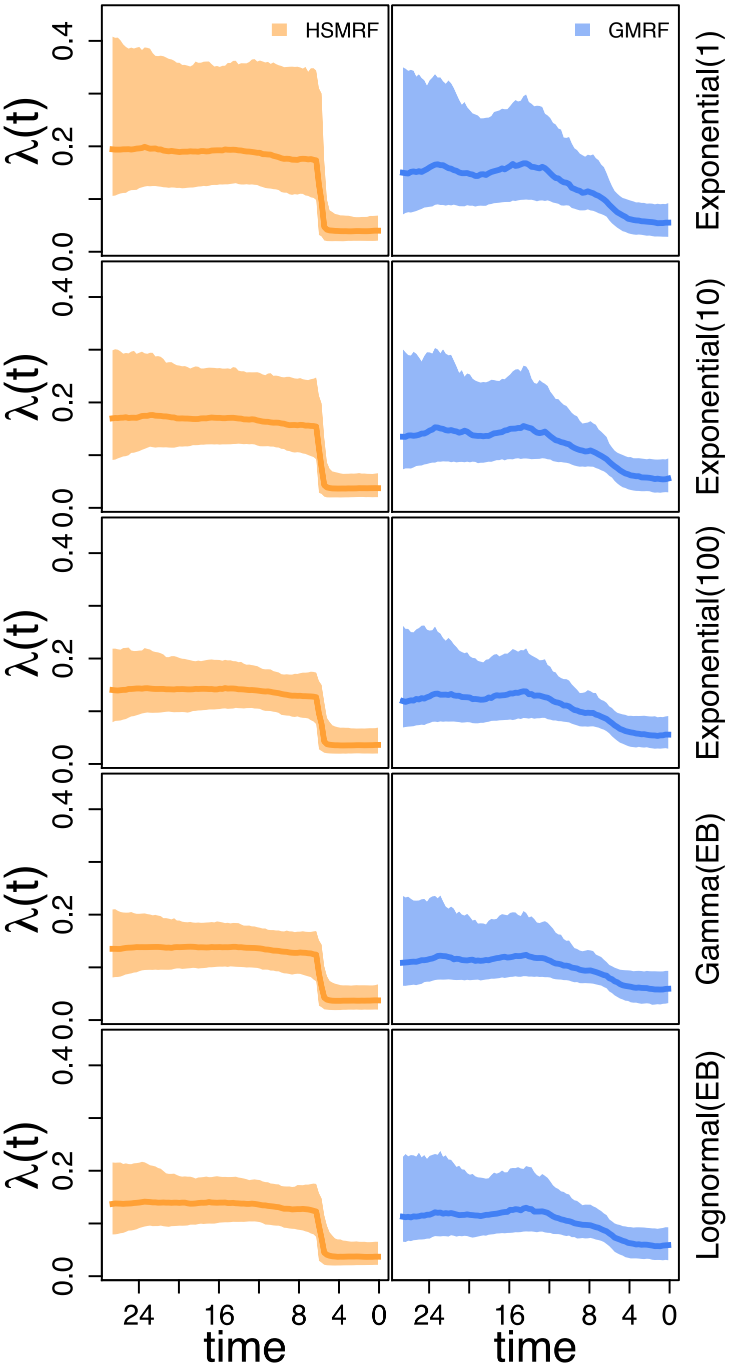


**Figure S21.** The effect of the death-rate prior on the estimated birth-rate trajectory. The tree is fixed to a tree from the posterior distribution from the Pygopodidae HSMRF analysis. The label on the right hand side is the prior on the death rate.

#### Setting the global shrinkage prior

As discussed in the main text, setting the halfCauchy(0,$\zeta$) prior on $\gamma$ is a crucial step in framing a random field model. Our strategy is inspired by the way Drummond and Suchard (2010) parameterize their random local clock model, but some adjustments have to be made because we work with completely continuous parameters. Namely, Drummond and Suchard (2010) choose a prior that places 50% probability on a model with 0 shifts, but for our models this implies a non-negligable probability of the process ending at a value over a one-trillion-fold changed from the initial. Because neither random field model is truly binary, even a process that shows no "rate shifts" can exhibit rather substantial variation throughout the history of the process. We therefore set our prior based on the expected number of shifts (the prior mean), and choosing this to be ln(2) aligns our mean with the mean of the prior in Drummond and Suchard (2010) [46].

There are three other classes of approaches to setting $\zeta$ that may be useful in different circumstances. In the first class of approaches, one commonly bounds the marginal variances of the GMRF, such that the probability that the marginal variance of the field at $t_i$ exceeds some reference value is $\alpha$ (which may be arbitrarily chosen to be 0.05) [47, 32]. This approach is not immediately amendable to parameterization via biological intuition, however. Alternately, it is possible to employ noisy estimators to bound the total variation of the random field [29]. This approach is applicable to modeling effective population sizes by using the skyline estimator [73] to get fast per-interval maximum likelihood estimates, then using the variances of these estimated effective population sizes to set $\zeta$. While there is no equivalent procedure available for birth-death processes, assuming a pure-birth process would allow the use of a skyline-like estimator to bound the process. However, such skyline-based approaches are only applicable in situations in which a tree already exists for the taxa; if there is no tree there is nothing from which we can estimate skyline birth rates. Therefore, while this approach can be quite useful, we prefer one that can be done before any trees have been estimated.

The third set of approaches considers each of the horseshoe distribution as producing variables that are binary, either effectively 0 or not. The application of horseshoe priors to regression models has been well studied in this framework [48]. The horseshoe introduces shrinkage weights, $\kappa_i$ on the model parameters $\beta_i$, such that at $\kappa_i = 1$ the posterior mean of $\beta_i$ is 0 and it is effectively not a model parameter (it is this distribution for which the horseshoe was named). Where analytical maximum likelihood estimators exist, it is possible to construct a prior on the effective number of parameters in the model. Given the complex dependencies among adjacent time-intervals in both birth-death processes and clock models, however, this approach seems out of reach.

#### Setting a prior for $\phi$

As with our prior on $\lambda_1$, it is possible to obtain an empirical estimate of the sampling rate $\phi$. Let $N(t)$ be the number of lineages alive at time $t$ (measuring time in time units after the origin, rather than our usual present to past). Assuming a constant rate model, taking $d = \lambda - \mu - \phi r$, and ignoring conditioning on the survival of the tree, the expected number of lineages at time $t$ is $\mathbb{E}(N(t)) = e^{d\times t}$ if we start with a single lineage. If we start with two lineages (as we do if we start at the MRCA), we have instead $\mathbb{E}(N(t)) = 2e^{d\times t}$. The rate of adding samples to the tree at time $t$ is $\phi N(t)$, with expectation $\phi e^{d\times t}$, or $2\phi e^{d\times t}$ if we start at the MRCA. Integrating, we get the expected number of serial samples up to time $t$ being $S(t_{or}) = \phi(e^{d\times t} - 1)/d$, or $S(t_{or}) = 2\phi(e^{d\times t} - 1)/d$ if we start at the MRCA. This means we can use the method of moments to obtain $\hat{\phi} = d \times S(t)/(e^{d\times t} - 1)$, or $\hat{\phi} = d \times S(t)/(2(e^{d\times t} - 1))$ if we start at the MRCA. This of course requires $d$, but as we have already discussed, estimation of $d$ is also possible by the method of moments. Our formula for $d$ in the serially sampled case can also be simplifed, since $S = B_{obs} + 2$, or $S = B_{obs} + 2$ if we start at the MRCA, meaning that in either case, we have $d =$ In practice one might wonder how well this approach performs, since we are repeatedly using the data to estimate these parameters. Using our heterochronously sampled simulations, we find that it works surprisingly well (Figure S22).

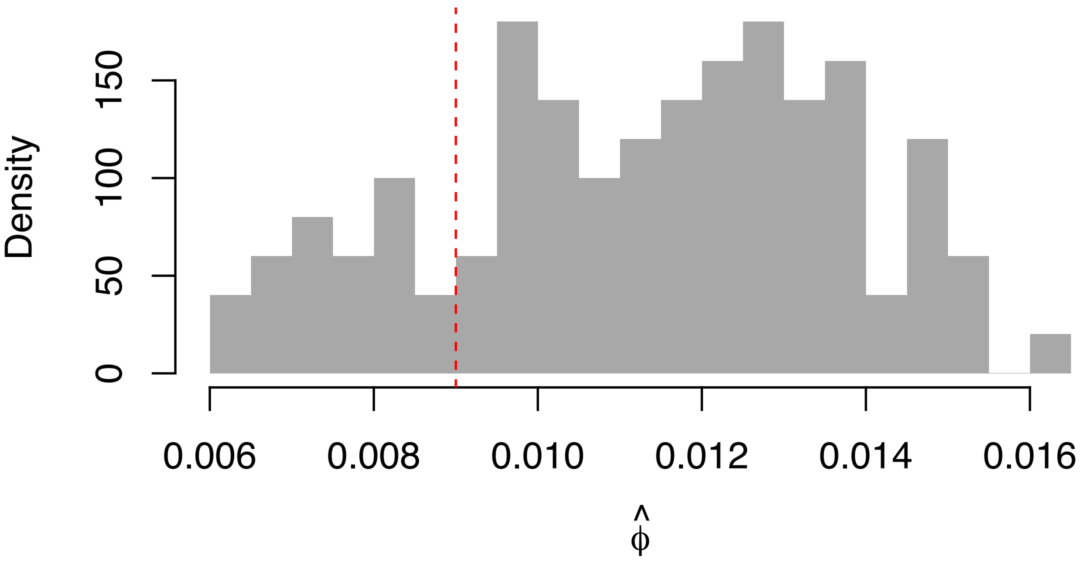


**Figure S22.** Performance of our proposed method of moments estimator $\hat{\phi}$ on our simulated heterochronously sampled trees. The true simulating value (0.009) is in red.

#### Diversification in Pygopodidae

We obtained the multiple sequence alignment for the entire dataset of Brennan and Oliver (2017) from their Dryad repository [74]. We extracted sequences for Pygopodidae manually in AliView [75], keeping only one representative for each species (the representative with the most sequence data), which left some gap-only sites that we later removed. Since our analyses concern a much smaller number of taxa, representing less variation at any site in the alignment, we used PartitionFinder 1.1.1 [76] to select subsets of the alignment for analysis. In PartitionFinder, we set branch lengths unlinked (such that different partitions should use independent molecular clock models), using BIC for model selection, and set the set of allowable substitution models to (JC, JC+G, JC+I, HKY, HKY+G, HKY+I, GTR, GTR+G, GTR+I), to avoid parameter identifiability issues with +I+G mixture models. The best schemes identified 2 subsets for Pygopodoidae, with the best substitution model being GTR+G for both [36].

In our analyses, for each subset we applied an uncorrelated lognormal clock model (UCLD) [35]. We placed an exponential(rate=3) prior on the log-scale standard deviation and a Normal(ln(0.001),4*0.587405) prior on the log-scale mean (yielding a prior median substitution rate of 0.001 per site per million years with a 95% prior CI of [1e-6,1]). Following Brennan and Oliver (2017), we applied uniform calibrations to the genera *Apprasia* (uniform(8.5,17)) and *Delma* uniform((14,22.5)) [6]. We used their node calibration for Pygopodidae (uniform(19.5,29.0)) as our root age prior. We ran 4 chains for 500,000 iterations for both the GMRF and HSMRF models (with 218 moves per iteration), downsampling to every 100th sample. This is the equivalent of 109 million generations per chain in a program like BEAST [59], sampled every 21,800.

We used the same rank-based PSRF procedure to assess convergence of the empirical analyses as for the simulated analyses. In convergence diagnostics, we ignore the branch-rate parameters as the recorded parameters in the log-file are not comparable across trees. These diagnostics revealed convergence issues with the substitution model for one of the HSMRF replicate analyses. While all other parameters had acceptable PSRF for the 4 HSMRF replicates, convergence is most properly addressed in an all-or-none framework, so we chose to exclude this entire chain from downstream analyses. Discarding this chain, the maximum rank-PSRF for the HSMRF analyses was 1.0008, and for the GMRF it was 1.0001. We were unable to calculate the effective sample size of the HSMRF chains directly due to some extremely large sampled values for the speciation rate. We thus chose to compute the effective sample size of the rank-transformed (but not folded) parameters for all chains, which should be correlated with (if not equal to) the effective sample size of the untransformed parameters. The rank-ESS of all parameters (pooled across all remaining chains) was above 2272 for all HSMRF parameters and above 2737 for all GMRF parameters.

#### HIV Dynamics in Russia and Ukraine

We obtained the multiple sequence alignment for the *env* dataset of Vasylyeva *et al.* from the first author [62]. Following previous analyses, we employed an unpartitioned GTR+G substitution model. We used a UCLD clock model, with a Normal(-9.21034,2.34962) prior on the log-mean (corresponding to a prior 95% CI of [1e-6,1e-2] substitutions per year) and an Exponential(3) prior on the clock log-SD. We used RAxML [77], using the GTRCAT substitution model, and TreeDater [78], with a strict clock, to obtain a starting tree for analysis and an estimate of the age of the tree. We used this estimate (29.1) to set a relatively diffuse Normal(29.1,5.0) prior on the tree age, with a truncation at 18.0, the relative age of the oldest sample. To set our prior on the serial sampling rate, we employ the empirical-Bayes strategy outlined above, resulting in a prior median sampling rate of 0.13. Back-of-the-envelope calculations of the sampling rate (using either the total number of infections or the numbers of samples out of the numbers of cases) suggest a rate on the order of $10^{-5}$. Future work may elaborate whether this discrepancy is common and what, if any, its effect is.

We ran 4 chains for 500,000 iterations for both the GMRF and HSMRF models (with 335 moves per iteration), downsampling to every 100th sample. This is the equivalent of 167.5 million generations per chain in a program like BEAST [59], sampled every 33,500. We performed convergence checks as with Pygopodidae, resulting in 2 GMRF-based model run failing and 2 HSMRF-based model runs failing. Discarding these chains, the maximum rank-PSRF for the HSMRF analyses was 1.003, and for the GMRF it was 1.003. The rank-ESS of all parameters (pooled across all remaining chains) was above 664 for all HSMRF parameters and above 317 for all GMRF parameters.

In Figure S23, we present posterior distributions of $\gamma$ for our analyses. As there are rapid changes evident, one might expect that $\gamma$, especially for the GMRF-based model, would need to be large to explain these changes. This is in fact what is seen: for both models, the posterior distributions of $\gamma$ are pulled up from the prior, and for the GMRF-based model the change is quite large. This also helps to explain the differences in the inferred $R_e(t)$ in the mid-1990s: since for the GMRF-based model $\gamma$ is so large, it infers substantial variability in this time, while the HSMRF-based model with a smaller $\gamma$ smooths out this variability.

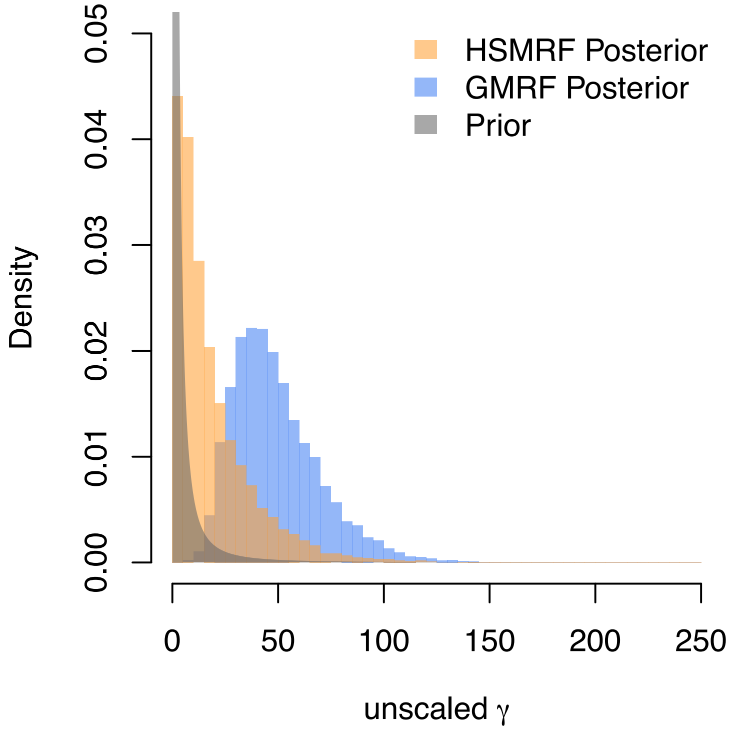


**Figure S23.** Comparison of the global scale parameter $\gamma$ for the HIV analysis. The grey curve is the halfCauchy(0,1) prior used on the unscaled $\gamma$, the orange histogram is the posterior distribution inferred by the HSMRF-based model, and the blue histogram the posterior distribution inferred by the GMRF-based model. The GMRF-based model requires a very large $\gamma$ in order to accomodate the periods of rapid change inferred, while the HSMRF-based model does not require such a large value.

#### Comparison to BEAST

For time-varying birth-death process models, the current state-of-the-art is the birth-death skyline model as implemented in the BEAST 2 [79] package BDSKY [17]. For serially-sampled datasets, this package implements a model in terms of the compound parameters $\boldsymbol{R_e} = \boldsymbol{\lambda}/(\boldsymbol{\mu}+\boldsymbol{\phi r})$, $\boldsymbol{\delta} = \boldsymbol{\mu}+\boldsymbol{\phi r}$, and $\boldsymbol{s} = (\boldsymbol{\phi r})/(\boldsymbol{\mu}+\boldsymbol{\phi r})$, though in all cases it is assumed that $r = 1$. These parameters are all assumed to be iid in some number of intervals that the user defines. To compare our results to the state-of-the-art, we used BEAST 2.6 [79] to perform joint inference of phylogeny and phylodynamic parameters. Comparability cannot be perfect, since some parameters of the model are different, however, we attempted to keep priors similar where possible and biologically motivated otherwise. Due to some difficulty with clock model convergence in preliminary analyses, we placed a tighter prior on the clock rate informed by our estimated clock rate from the `RevBayes` analyses. Our prior, a Lognormal(-4.9,0.587), corresponds to the clock rate being within approximately one order of magnitude of the clock rate estimated in our `RevBayes` analyses We assumed a single rate of becoming noninfectious, $\delta$, and a single sampling proportion, $s$, analogous to our assumptions that $\mu$ and $\phi$ are constant. We placed a Lognormal(-2.272,0.073) prior on $\delta$ and a Beta(1,19) prior on $s$, reflecting our knowledge about the rate of becoming noninfectious (absent treatment) and the fact that we have a small sample out of a much larger epidemic. We placed Lognormal(0.0,0.821) priors on $R_{ei}$, reflecting that the effective reproductive number should likely be lower than 5, and the possibility that $R_e$ may dip below 1.0. We employ a grid with 15 intervals, with break-points specified to occur at 1990,1991,...,2002,2003. Given that there are few infections inferred to have occurred post-2003 or prior to 1990 based on our analyses using the GMRF-based and HSMRF-based models, our most recent grid cell begins in 2011 and ends in 2003, while our oldest grid cell contains all times prior to 1990. Convergence diagnostics showed a maximum rank-based PSRF of 1.001 and a minimum rank-based ESS of 4932.

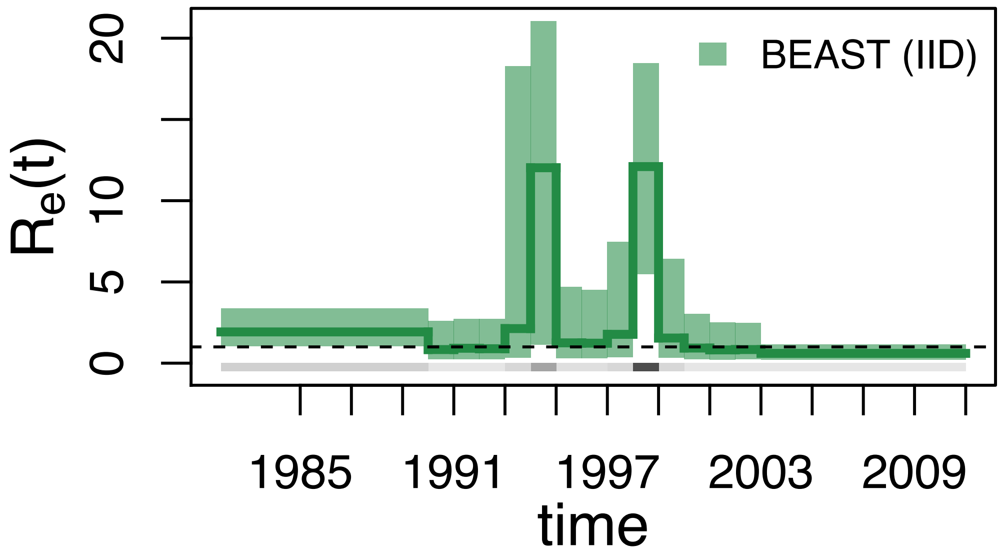


**Figure S24.** Analyses of the HIV dataset with BEAST. Plotted are posterior median trajectories (dark lines) and 90% credible intervals (shaded regions). Time is plotted as calendar time. A line at $R_e = 1$ is provided for convenience, as below this threshold the epidemic cannot be sustained. In grey is a heatmap of the inferred divergence times.

The results of our BEAST analysis bear many similarities to the results from our GMRF-based model. Qualitatively, the pattern is similar, two peaks in $R_e(t)$ at approximately 1994 and 1998, each proceeded by a decrease, a low rate at the present, and a higher rate in the past. The 2ln Bayes Factor for a decrease at the end of the 1990s is 6.36 (strong support), while the 2lnBF in favor of an increase at the beginning of the decade is 6.09 (strong support). However, where the GMRF shows clear signs of an elevated rate for the entirety of the 1990s, the BEAST analysis shows a very strong decrease in the middle of the decade. The 2lnBF for the decrease after 1995 is 4.62 (positive support) and for the increase before 1998 is 5.79 (positive support). The inferred rates in the peak periods are implausibly high, with the estimated $R_e(t)$ at the peaks being much higher than in our HSMRF-based and GMRF-based model analyses at approximately 12.

The results from BEAST form one end of the spectrum of temporal smoothing, with the GMRF-based model in the middle and the HSMRF-based model on the opposite side. When all intervals are iid, there is no smoothing and thus no pooling of information among adjacent (or close) intervals, and any apparent change in the birth rate will be picked up. When intervals are temporally autocorrelated, these apparent changes are weighted against a prior belief that the rate is likely similar over small periods of time, requiring more evidence before a shift is inferred. The HSMRF-based model infers $R_e(t) > 1$ prior to 1991, where both the less-smoothed GMRF-based model and unsmoothed BEAST analysis suggest $R_e(t) < 1$ prior to the first increase. The HSMRF-based model infers a high $R_e(t)$ throughout most of the 1990s, while the GMRF-based model infers two peaks of $R_e(t)$ with a notable decrease between them, and the BEAST analysis exaggerates this peak (and shows a posterior probability of 0.3 to 0.4 that $R_e(t) < 1$ in this range). In the case of the HIV results, smoothing and sharing information across intervals produces analyses more concordant with other lines of epidemiological research (which do not show these decreases).

#### Marginal likelihoods and Metropolis Coupled MCMC

As our MCMC sampler is somewhat atypical in phylogenetics, and may thus be unfamiliar to readers, we now examine how the sampler interacts with common variants of MCMC in phylogenetics. Calculating marginal likelihoods via path sampling [80] or stepping stone [81] both require running a number of MCMC chains, each of which raises the likelihood of the model to an increasing power $0 \leq \beta \leq 1$. Metropolis-coupling employs multiple MCMC chains where all but one (the so-called cold chain) have their target posteriors raised to a power $0 \leq \beta < 1$. While these are not issues for the standard Metropolis-Hastings moves in most phylogenetic inference software, they can pose problems for other types of samplers such as Gibbs samplers. Employing a Gibbs Sampler requires that the conditional distributions of the parameters being sampled remain unchanged and cannot be raised to any power. This means that the distribution on all parameters above and those directly below $\boldsymbol{\sigma}$ and $\gamma$ in the model DAG (see Figure S18) cannot be raised to any power $\beta \neq 1$. The elliptical slice sampler requires that the prior on $\boldsymbol{\Delta}$ be multivariate normal, so nothing above $\boldsymbol{\Delta}$ in the model DAG can be raised to a power. Taken together, these imply that anything below $\boldsymbol{\Delta}$ can be raised to a power, and thus marginal likelihoods may be computed, but that these samplers are incompatible with Metropolis-coupled MCMC. For both Gibbs samplers and for the elliptical slice sampler, `RevBayes` will throw an error if the user attempts to raise any distributions to unallowed powers.

#### Implementation details

In `RevBayes`, a distinction is made between the EBD, which has no serial sampling, and the episodic birth-death sampling treatment process, which allows for serial sampling and conditional death upon sampling (treatment). That is, the EBD is the model of Stadler (2011) and Höhna (2015), whereas the EBDSTP is the model of Gavryushkina *et al.* (2014) [15, 16, 18]. When there is no serial sampling in the model ($\phi(t) = 0$) and there are tips only at the present ($\boldsymbol{t}_{\boldsymbol{\Phi}} = t_{\Phi} = 0$), the models are eactly equivalent. In these cases, we employ the simpler EBD model in our Rev scripts, though we note that changing back is a simple matter of replacing calls to `dnEBDP` with calls to `dnBDSTP` in `RevBayes`.
